# Structural basis for the unique multifaceted interaction of DPPA3 with the UHRF1 PHD finger

**DOI:** 10.1101/2022.06.14.496052

**Authors:** Keiichi Hata, Naohiro Kobayashi, Keita Sugimura, Weihua Qin, Deis Haxholli, Yoshie Chiba, Sae Yoshimi, Gosuke Hayashi, Hiroki Onoda, Takahisa Ikegami, Christopher B. Mulholland, Atsuya Nishiyama, Makoto Nakanishi, Heinrich Leonhardt, Tsuyoshi Konuma, Kyohei Arita

## Abstract

Ubiquitin-like with PHD and RING finger domain-containing protein 1 (UHRF1)-dependent DNA methylation is essential for maintaining cell fate during cell proliferation. Developmental pluripotency-associated 3 (DPPA3) is an intrinsically disordered protein that specifically interacts with UHRF1 and promotes passive DNA demethylation by inhibiting UHRF1 chromatin localization. However, the molecular basis of how DPPA3 interacts with and inhibits UHRF1 remains unclear. We aimed to determine the structure of the mouse UHRF1 plant homeodomain (PHD) complexed with DPPA3 using nuclear magnetic resonance. Induced α- helices in DPPA3 upon binding of UHRF1 PHD contribute to stable complex formation with multifaceted interactions, unlike canonical ligand proteins of the PHD domain. Mutations in the binding interface and unfolding of the DPPA3 helical structure inhibited binding to UHRF1 and its chromatin localization. Our results provide structural insights into the mechanism and specificity underlying the inhibition of UHRF1 by DPPA3.

## INTRODUCTION

Cytosine DNA methylation of the CpG sequence in mammals plays a pivotal role in embryogenesis, retrotransposon silencing, X-chromosome inactivation, genome imprinting, and carcinogenesis(1). Mammalian cells undergo two waves of methylation changes: DNA methylation and demethylation(2). After fertilization, DNA methylation patterns derived from gametes are erased during early embryogenesis and re-established during cellular development(3). DNA methylation patterns in primordial germ cells (PGCs) are erased globally, and sex-specific DNA methylation patterns are established during germ cell development(2, 4). After establishment, cell-type-specific DNA methylation patterns are faithfully propagated after each cycle of replication to maintain cellular identity. DNMT1, a maintenance DNA methyltransferase, and UHRF1 (ubiquitin-like PHD and RING finger domain-containing protein 1, also known as Np95/ICBP90), which is a ubiquitin E3-ligase and recruiter of DNMT1, are essential for DNA methylation maintenance(5–8). UHRF1 specifically binds to hemi-methylated DNA and ubiquitinates histone H3 and PCNA-associated factor 15 (PAF15) to recruit DNMT1 to chromatin and replication sites(9–13). The two distinct ubiquitin signals are involved in replication-coupled and uncoupled DNA methylation maintenance(9, 14, 15).

UHRF1 functions as a reader of epigenetic marks, hemi-methylated DNA, and the H3K9me2/3 modification(16–19) to regulate its ubiquitination activity(20). Further, it serves as a binding platform for DNA replication factors and epigenetic modifiers, which are involved in DNA methylation maintenance, gene expression, DNA damage repair, and tumorigenesis(9, 21–28). Additionally, mouse DPPA3 (developmental pluripotency associated 3, also known as Stella/PGC7; hereafter mDPPA3) is a novel ligand that regulates the binding of UHRF1 to chromatin(29, 30). mDPPA3, an intrinsically disordered protein (IDP), is specifically expressed in PGCs, oocytes, and preimplantation embryos, and plays an important role in the formation of oocyte-specific DNA methylation patterns by preventing excessive *de novo* DNA methylation mediated by UHRF1(29, 31–33). Overexpression of mDPPA3 in somatic cells such as NIH3T3, HEK293, and mouse embryonic stem cells, and in non-mammalian species, results in genome-wide DNA demethylation, indicating that mDPPA3 is a passive DNA demethylation factor that inhibits the cellular functions of UHRF1(29, 34, 35).

UHRF1 has five functional domains: a ubiquitin-like domain (UBL), tandem tudor domain (TTD), plant homeodomain (PHD) finger, SET and RING associated domain (SRA), and really interesting new gene (RING) (Figure 1A). mDPPA3 interacts with the UHRF1 PHD finger, resulting in the inhibition of chromatin binding of UHRF1(35, 36). The UHRF1 PHD finger also interacts with the N-terminal ^1^ARTK^4^ in histone H3 (H3) and ^1^VRTK^4^ in PAF15, in which strict recognition of the main chain amino group at the first residue of H3 and PAF15 is critical for their ubiquitination(9, 19). The PHD finger is one of the largest families of chromatin-reader domains. They have been found in more than 100 human proteins, most of which recognize K4 methylation state in the H3 N-terminal tail(37). Notably, with a few exceptions, the recognition mode of the first amino acid residue amino group in ligands is conserved among almost all PHD fingers(38). Indeed, acetylation of the N-terminus of the H3 tail abolished binding to the PHD finger of UHRF1(19). Given that the ARTK/VRTK-like sequence is not present at the N-terminus of mDPPA3, the molecular mechanisms by which the UHRF1 PHD finger recognizes DPPA3 is unknown.

**Figure 1:**
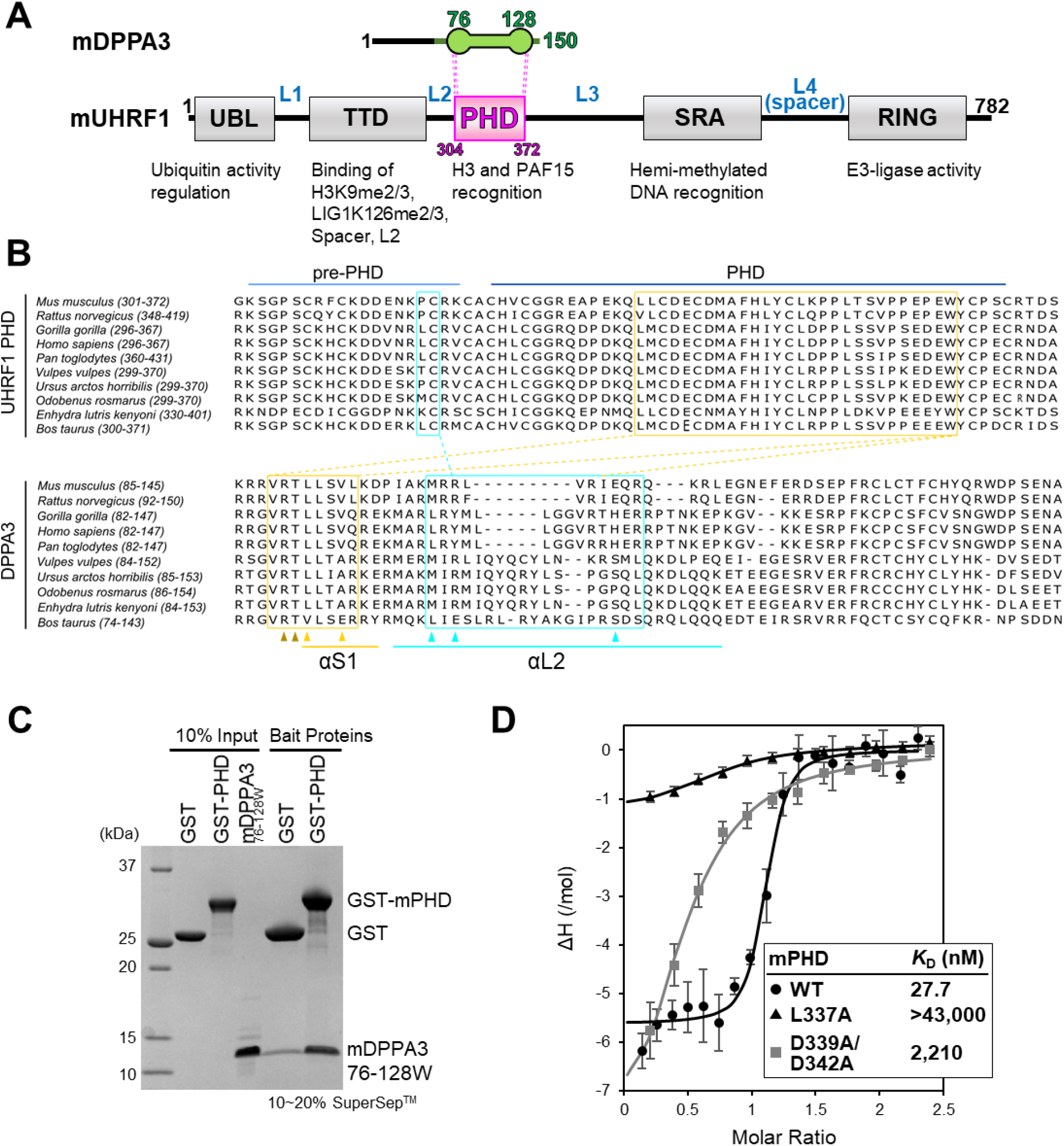
Characterization of the interaction between mUHRF1 and mDPPA3. **(A)** Schematic of the domain organization of mouse UHRF1 and DPPA3. L1-L4 indicate the connecting linkers. **(B)** Protein sequence alignment of the UHRF1 pre-PHD/core-PHD domains and DPPA3 residues 85-127 between different species with Jalview software. The cyan and yellow dotted lines indicate the interaction interface identified in this study. **(C)** GST pull-down experiments using mPHD and mDPPA3_76-128W_. Bait proteins mean GST-fused mUHRF1 that is immobilized on GST-beads. Proteins are stained by Coomassie Brilliant Blue. **(D)** ITC measurements for the mPHD wild-type (WT)/mutant and mDPPA3_76-128W_. Superimposition of enthalpy change plots with standard deviations.

Here, we aimed to determine the solution structure of mouse UHRF1 PHD (mPHD) in complex with the C-terminal fragment of mDPPA3 using nuclear magnetic resonance (NMR), and identified the unique multifaceted interaction of mDPPA3 with mPHD. Although the ^88^VRT^90^ cassette of mDPPA3 is not located at its N-terminus, unlike H3 or PAF15, we found that it was recognized by a shallow acidic groove on the mPHD in a manner similar to the N- terminus of H3 and PAF15. Structural induction of the two α-helices of mDPPA3 provided several additional binding sites for mPHD, which plays an important role in stable complex formation. Structure-guided mutagenesis and functional assays using *Xenopus* egg extracts and mouse embryonic stem cells (mESCs) helped evaluate the key amino acid residues of mDPPA3 that negatively regulate the binding of mUHRF1 to chromatin, and shed light on the mechanisms underlying chromatin delocalization of mUHRF1 by mDPPA3. Our data provide insight into the diversity of recognition of ligand proteins by the PHD finger and contribute to the understanding of its key role in epigenetic maintenance.

## MATERIALS AND METHODS

### Protein expression and purification

For NMR and ITC experiments, cDNA of mouse UHRF1 PHD (residues 304–372) was sub-cloned into a pGEX6P-1 plasmid (Cytiva) at 5’-BamHI and 3’-XhoI sites for protein expression with N-terminal glutathione S-transferase (GST). The protein was expressed in *E*.*coli* BL21 (DE3) in Luria–Bertani medium (LB) containing 50 μg/mL ampicillin. When the optical density at 660 nm (O.D._660_) of the cells reached 0.7, 0.2 mM isopropyl β-d- thiogalactoside (IPTG) was added to the medium and the cells were further harvested for 3 h at 37°C. The cells were suspended with lysis buffer (40 mM Tris-HCl [pH 8.0], 300 mM NaCl, 10% glycerol, 30 μM Zn-acetate, 0.5 mM tris(2-carboxyethyl)phosphine (TCEP)) and disrupted by sonication. After the cell debris were removed by centrifugation, the supernatant was loaded onto a GST-affinity column, GS4B (Cytive). After the protein was eluted from the column using reduced glutathione, the GST-tag was cleaved by HRV-3C protease. The sample was further purified using HiTrap Q HP anion-exchange chromatography (Cytiva). Final purification was performed using HiLoad 26/600 Superdex 75 size-exclusion chromatography (Cytiva) equilibrated with buffer 1×phosphate-buffered saline (adjusted to pH 7.0; PBS) containing 1 mM DTT.

mDPPA3, residues 76–128W, for NMR and ITC experiments was expressed as a six histidine-tagged ubiquitin fusion protein. The procedures for cell culture were the same as that for mPHD. The cells were suspended in lysis buffer (30 mM HEPES [pH7.5], 400 mM NaCl, 0.1% Nonidet P-40 (NP-40), 40 mM imidazole). After cell lysis by sonication and removal of cell debris by centrifugation, the supernatant was loaded to histidine-tag affinity column Ni Sepharose 6 Fast Flow (Cytiva), and the sample was eluted from the column using an elution buffer containing 500 mM imidazole. Next, the histidine-tag was removed by *Saccharomyces cerevisiae* ubiquitin carboxyl-terminal hydrolase YUH1. The sample was further purified using HiTrap SP HP cation-exchange chromatography (Cytiva) and finally purified by HiLoad 26/600 Superdex 75 size-exclusion chromatography equilibrated with 1×PBS buffer containing 1 mM DTT.

For structure determination using NMR, cDNA of mDPPA3 (residues 76–127) was inserted into the 3’ end of the mPHD (residues 304–372) in pGEX6P-1 plasmid for expression as fusion protein. The procedures for cell culture of the mPHD-mDPPA3 were same as for mPHD. The cells were suspended with lysis buffer (40 mM Tris-HCl [pH 8.0], 300 mM NaCl, 10% glycerol, 30 μM Zn-acetate, 0.5 mM TCEP) and then disrupted by sonication. After the cell debris was removed by centrifugation, the supernatant was loaded onto GS4B. The protein was eluted from the column using reduced glutathione, and then GST-tag was cleaved by HRV- 3C protease. The sample was further purified using HiLoad 26/600 Superdex 75 size-exclusion chromatography equilibrated with 1×PBS buffer containing 1 mM DTT.

For preparation of ^15^N-labeled or ^13^C, ^15^N-double labeled mPHD, mDPPA3 and mPHD-mDPPA3, M9 minimal media containing 0.5 g/L ^15^NH_4_Cl or 0.5 g/L ^15^NH_4_Cl and 1 g/L ^13^C-glucose was used instead of LB media. Site-directed mutagenesis was performed by designing two primers containing the mutations. The mutants of mPHD and mDPPA3 were purified using the same protocol.

### GST pull-down assay

MagneGST™ Glutathione Particles (Promega) were used for the assay. The truncated GST-mUHRF1 (10 μg) were immobilized on the beads (10 μL) equilibrated with the binding buffer (20 mM Tris-HCl [pH 7.5], 150 mM NaCl, 1 mM DTT, 10% glycerol, and 0.5 % NP-40). After the unbound proteins were washed-out thrice by the 200 μL of the binding buffer, 10 μg of C-terminal fragment of mDPPA3 (residues 61–150 or 76–128W) was incubated with the beads for 2 h at 4°C. After incubation, the unbound protein was washed thrice using 200 μL of the binding buffer. The bound proteins were boiled for 2 min at 95°C in an SDS sample buffer and analyzed by SDS-PAGE.

### ITC

Microcal PEAQ-ITC (Malvern) was used for the ITC measurements. Wild-type and mutants of mPHD and mDPPA3 were dissolved in 10 mM HEPES (pH 7.5) buffer containing 150 mM NaCl and 0.25 mM TCEP. All measurements were carried out at 293K. The data were analyzed with Microcal PEAQ-ITC analysis software using a one-site model. For each interaction, at least three independent titration experiments were performed to show the dissociation constants with the mean standard deviations.

### NMR

All NMR experiments were performed on Bruker BioSpin Avance III HD spectrometers with a TCI triple-resonance cryogenic probe-head with basic ^1^H resonance frequencies of 500.13, 800.23 and 950.15 MHz. Three-dimensional (3D) spectra for main-chain signal assignments: HNCACB, HN(CO)CACB, HNCA, HN(CO)CA, HNCO and HN(CA)CO, and for side-chain signal assignments: HBHA(CBCACO)NH, H(CCO)NH, CC(CO)NH, HCCH-TOCSY and (H)CCH-TOCSY, for structure analysis: ^1^H-^13^C NOESY- HSQC, and ^1^H-^15^N NOESY-HSQC spectra were acquired at 293 K for 660 μM [^13^C, ^15^N]- mPHD-mDPPA3 fusion protein dissolved in PBS buffer (pH 7.0) containing 1 mM DTT and 5% D_2_O. For the main-chain signal assignments of isolated proteins, 500 μM [^13^C, ^15^N]-mPHD in the free, 350 μM [^13^C, ^15^N]-mPHD in the complex with mDPPA3 at molar ratio of 1:2, 500 μM [^13^C, ^15^N]-mDPPA3 in the free state, and 350 μM [^13^C, ^15^N]-mDPPA3 in the complex with mPHD at molar ratio of 1:2 were used in the buffer same as the fusion protein. The spectral widths (the number of total data points) of each spectrum were 24 ppm (2,048) for the ^1^H dimension, and 30 ppm (256) for the ^15^N dimension. All 3D spectra except for ^1^H-^13^C NOESY- HSQC were acquired by means of non-uniform sampling (NUS) to randomly reduce t1 and t2 time-domain data points typically around 25%. The uniformly sampled data were reconstructed from the raw NMR data according to the sparse sampling schedules using several techniques such as IST, SMILE, MDD and IRLS(39–41). Chemical shift perturbation experiments were performed by recording 2D ^1^H-^15^N HSQC spectra of 30 μM [^15^N]-mPHD dissolved in the same buffer. All NMR spectra were processed using NMRPipe(42). For the NMR analysis, an integrated package of NMR tools named MagRO-NMRViewJ, version 2.01.39 [the upgraded version of Kujira(43)], on NMRView(44) was used for automated signal identification and noise filtration using convolutional neural networks [CNNfilter(45)]. The filtered signal lists were applied to calculations for automated signal assignments by FLYA(46) and then the signal assignments were used for prediction of dihedral angles by TALOS+(47), and automated NOE assignments and structure calculation by CYANA(48) (Table 1). Finally, water refinement calculations by AMBER12 were performed for the lowest energy structures (20 models) (Table 2).

**Table 1.**
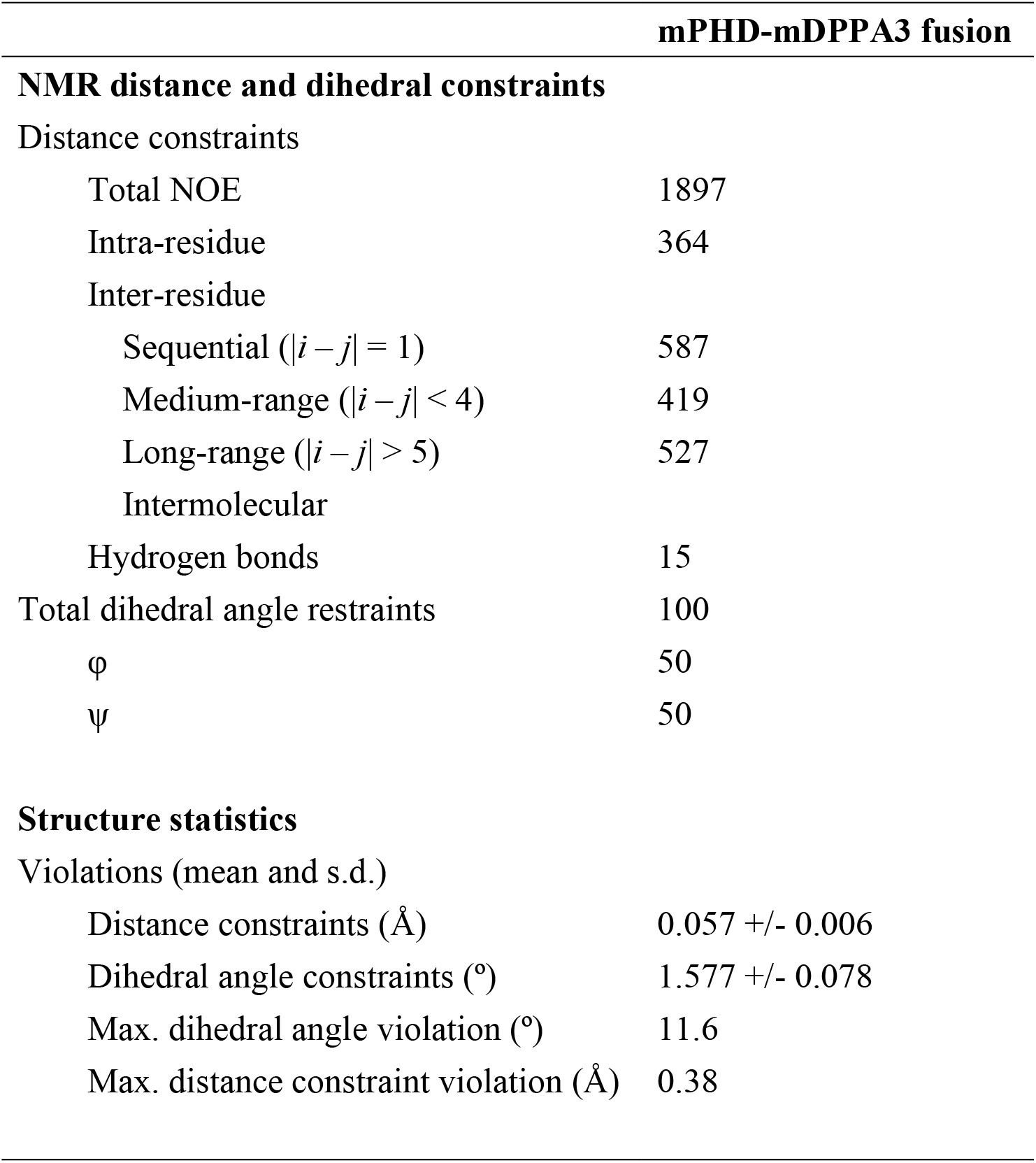
NMR and refinement statistics for protein structures.

**Table 2.**
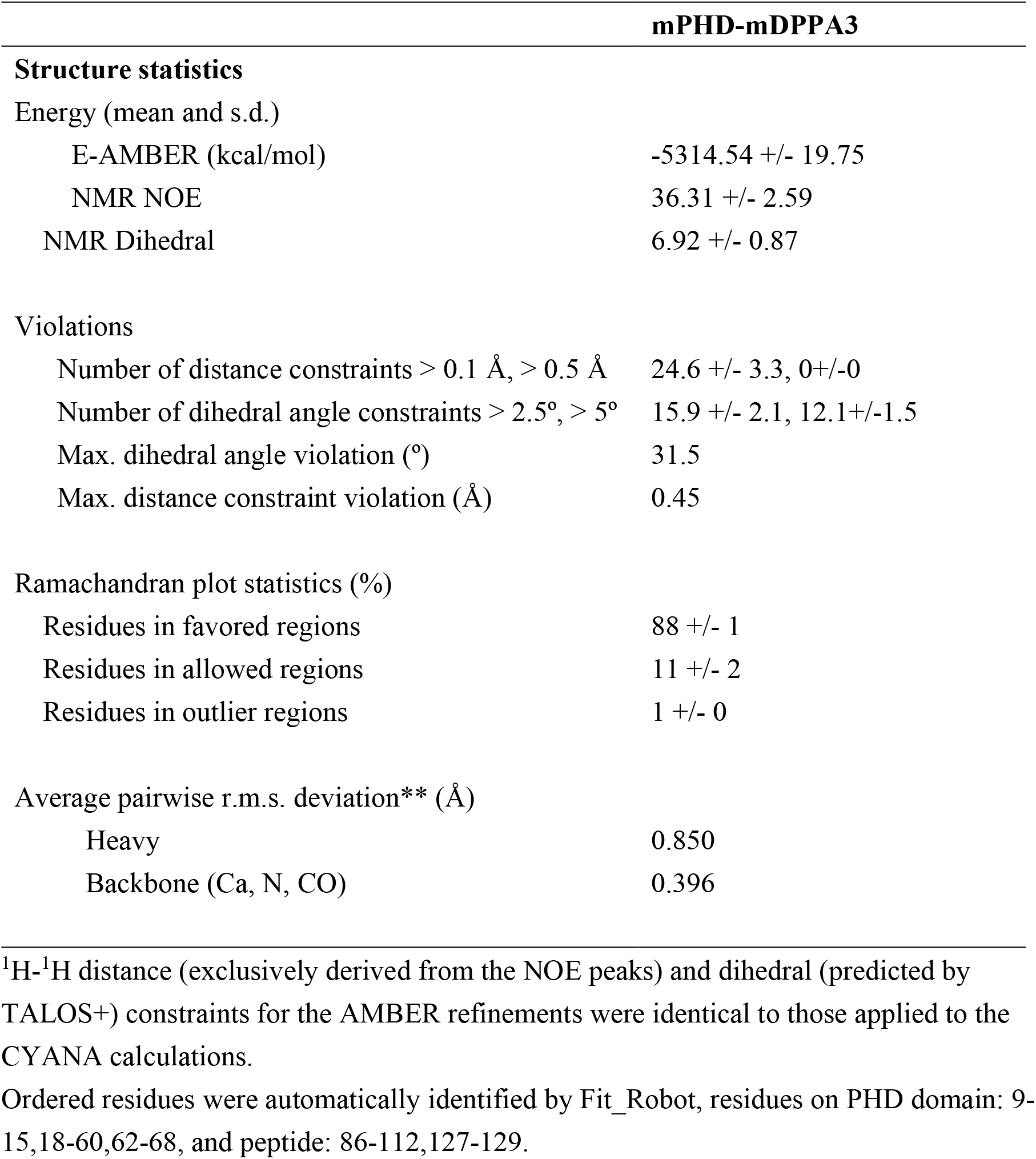
AMBER refinement statistics for complexes.

Furthermore, we ran fully automated peak picking and noise filtration on MagRO for all spectra required for FLYA calculation using structure calculation mode with peak tables for the spectra: ^1^H-^15^N HSQC, HNCO, HN(CA)CO, HNCA, HN(CO)CA, CBCA(CO)NH, HNCACB, HCCH-TOCSY,^15^N-edited NOESY, and ^13^C-edited NOESY. After the 1st FLYA calculation, ∼80% of backbone and ∼60% side-chain signals were automatically assigned. These assigned chemical shifts were imported into MagRO from the output flya.tab file, according to the following criteria: among 20 assigned chemical shift tables described in flya.tab, a cut-off value of 80% was set to eliminate poorly populated assigned chemical shifts after the final consolidation stage, and several proton signals such as Thr-OH, Ser-OH, His-H_e2_, His-H_d1_, and Phe-H_z_ were also neglected. Following visual inspection of the assigned data using 3D spectra such as H(CCCO)NH, CC(CO)NH and HBHA(CO)NH to confirm and correct the assignments, 2^nd^ FLYA calculation was performed with the confirmed assigned chemical shifts to assign the remaining signals. TALOS+ calculation was performed using chemical shifts for ^1^HN, ^15^N, ^13^C_a_, ^13^C_b_, and ^13^CO to predict the dihedral angles (phi and psi) of backbone. MagRO automatically converted the predicted dihedral angles to talosp.aco restrain file in the CYANA format except for the data which may not be trustful: low predicted order parameter (less than 0.8) and worse class annotated as “Warn” or “Dyn”, as well as with setting a minimum angle (±20-deg). In the CYANA calculation, we applied 6 upper- and lower-limit distance constraints (total 82) to form tetrahedral coordinates for each Zn^2+^ atom, which was topologically linked with pseudo residues “PL” and “LL2”. We assumed that the members of residues involved in the three Zn-finger coordinates from the typical fashion found in PHD domains, namely site1: C-C-C-C, site2: C-C-C-C, and site3: C-C-H-C.

For the obtained CYANA structures (20 models) with the lowest target function, implicit water refinement calculations were performed by AMBER12. Dihedral angle constraints (derived from TALOS+ prediction) and distance constraints including additional chirality and backbone omega angle constraints were converted for AMBER format using the SANDER tool(49). The distance constraints exported by CYANA (final.upl) were exclusively used (purely derived from experimental NOE data). Metal coordinate parameters, ZAFF.prep and ZAFF.frcmod, were appended to the standard force field, ff99SB, for the three zinc-finger cites according to the AMBER tutorial [http://ambermd.org/tutorials/advanced/tutorial20/ZAFF.htm]. In each initial stage of the refinement, 500 steps of energy minimization (250 steps of steepest gradient, followed by 250 steps of conjugate gradient decent) without electro-static energy and NMR constrain terms. The calculation was followed by a short molecular dynamics calculation (total 100 psec, time step 1.0 fsec, using SHAKE algorism for bond and angle stabilization) using electro-static energy based on generalized born model (salt concentration: 100 mM, disabled Surface Accessibility (SA) function, electro-static potential radius cut-off: 18 Å) and NMR constrain terms. After the temperature was gradually increased from 0 to 300 K for 1,500 steps, dynamic calculation at 300 K was ran for remaining steps. In the final stage of the refinement, 2,000 steps of energy minimization were performed with the same energy terms.

^1^H-^15^N heteronuclear NOE (het-NOE) spectra of mPHD-mDPPA3 were measured with and without ^1^H saturation applied before the start of the measurement(50). The het-NOE values were calculated as ratios of the signal intensities of the two spectra. For signal intensity of each residue, the experimental error was estimated from the target signal around noise amplitude.

For the competition experiments, ^1^H-^15^N HSQC spectra were measured at 293K for 80 μM [^15^N]-mPHD in the presence of mDPPA3_76-128W_ and/or the H3 1-37W peptide at molar ratios (mPHD:mPDDA3:H3) of 1:0:0, 1:2:0, 1:0:2 and 1:2:8.

### CD

Far-UV circular dichroism (CD) spectra were obtained using a JASCO J-720W model spectrometer. All samples were prepared to a concentration of 20 μM. Measurements were performed at 293K with a path length of 1 mm.

#### *In vitro* ubiquitination assay

Protein expression in *E. coli* and purification of mouse UBA1 (E1), human UBE2D3 (E2), mouse UHRF1 (E3), C-terminal FLAG tagged-H3_1-37W_ and ubiquitin were performed according to the methodology in previous reports(9). Ubiquitination reaction mixtures contained 100 μM ubiquitin, 200 nM E1, 8 μM E2, 3 μM E3, 5 mM ATP, and 10 μM C-terminal FLAG tagged-H3_1-37W_ in the presence and absence of 10 μM mDPPA3_76-128W_ in ubiquitination reaction buffer (50 mM Tris-HCl [pH 8.0], 50 mM NaCl, 5 mM MgCl_2_, 0.1% Triton X-100, 2 mM DTT). The mixture was incubated at 30°C for 3 h, and the reaction was stopped by adding 3×SDS loading buffer. The reaction was analyzed by SDS-PAGE, followed by Western blotting using 1/5,000 diluted anti-FLAG antibody (Cell Signaling Technology, #2368).

#### *Xenopus* interphase egg extracts and purification of chromatin

*Xenopus laevis* was purchased from Kato-S Kagaku and handled according to the animal care regulations at the University of Tokyo. Interphase egg extracts were prepared as described previously(11). Unfertilized *Xenopus* eggs were dejellied in 2.5% thioglycolic acid-NaOH (pH 8.2) and were washed three times in 0.2×MMR buffer (5 mM HEPES-KOH [pH 7.6], 100 mM NaCl, 2 mM KCl, 0.1 mM EDTA, 1 mM MgCl_2_, 2 mM CaCl_2_). After activation in 1×MMR supplemented with 0.3 μg/mL calcium ionophore, eggs were washed four times with EB buffer (10 mM HEPES-KOH [pH 7.7], 100 mM KCl, 0.1 mM CaCl_2_, 1 mM MgCl_2_, 50 mM sucrose). Packed eggs were crushed by centrifugation (BECKMAN, Avanti J-E, JS13.1 swinging rotor) for 20 min at 18,973×g. Egg extracts were supplemented with 50 μg/mL cycloheximide, 20 μg/mL cytochalasin B, 1 mM DTT, 2 μg/mL aprotinin, and 50 μg/mL leupeptin and clarified for 20 min at 48,400×g (Hitachi, CP100NX, P55ST2 swinging rotor). The cytoplasmic extracts were aliquoted and stored at -80°C. Chromatin purification after incubation in egg extracts was performed as previously described with modifications. Sperm nuclei were incubated in egg extracts supplemented with an ATP regeneration system (20 mM phosphocreatine, 4 mM ATP, 5 μg/mL creatine phosphokinase) at 3000–4000 nuclei/μL at 22°C. Aliquots (15 μL) were diluted with 150-200 μL chromatin purification buffer (CPB; 50 mM KCl, 5 mM MgCl_2_, 20 mM HEPES-KOH [pH 7.7]) containing 0.1% NP-40, 2% sucrose and 2 mM NEM. After incubating on ice for 5 min, extracts were layered over 1.5 mL of CPB containing 30% sucrose and centrifuged at 15,000×g for 10 min at 4°C. Chromatin pellets were resuspended in 1×Laemmli sample buffer, heated for 5 min and analyzed by SDS-PAGE. Recombinant Flag-tagged mDPPA3 was added to egg extracts at 400 nM.

DNA methylation was monitored by the incorporation of S-[methyl-3^H^]-adenosyl-L-methionine. Extracts supplemented with S-[methyl-3^H^]-adenosyl-L-methionine and sperm nuclei were incubated at 22°C in the presence or absence of recombinant Flag-tagged mDPPA3 protein. After 90 min incubation, the reaction was stopped by the addition of CPB containing 2% sucrose up to 300 μL. Genomic DNA was purified using a Wizard Genomic DNA Purification Kit (Promega) according to the manufacturer’s instructions. The incorporation of radioactivity was quantified with a liquid scintillation counter.

#### GST pull-down assay using *Xenopus* egg extracts

For GST pull-down experiments using *Xenopus* egg extracts, mouse DPPA3 cDNA was sub-cloned into a pGEX4T-3 plasmid using In-Fusion (Clontech) according to the manufacturer’s instructions. GST or GST-mDPPA3 proteins were expressed in *E*.*coli* BL21 (BL21-CodonPlus) by the addition of 0.1 mM IPTG to media followed by incubation for 12 h at 20°C. Bacteria cells were harvested and resuspended in lysis buffer (20 mM HEPES-KOH [pH 7.6], 500 mM NaCl, 0.5 mM EDTA, 10% glycerol, 1 mM DTT) supplemented with 0.5% NP-40 and protease inhibitors and were then disrupted by sonication on ice. After the cell debris was removed by centrifugation, the recombinant proteins were immobilized on GS4B resin by incubation for 2 h at 4°C. After the unbound proteins was washed-out by the lysis buffer, interphase egg extracts were incubated with the beads for 2 h at 4°C. After incubation, the beads were washed four times with CPB containing 2% sucrose, 500 mM KCl and 0.1% Triton X-100. The washed beads were resuspended in 20 μL of 2×Laemmli sample buffer and 20 μL of 1×Laemmli sample buffer, boiled for 5 min at 100°C, and analyzed by immunoblotting.

For FLAG-tagged protein expression in insect cells, 3×FLAG-tagged mDppa3 WT or mutants were sub-cloned into the pVL1392 vector. Baculoviruses were produced using a BD BaculoGold Transfection Kit and a BestBac Transfection Kit (BD Biosciences), following the manufacturer’s protocol. Proteins were expressed in Sf9 insect cells by infection with viruses expressing 3×FLAG-tagged mDPPA3 WT or its mutants for 72 h at 27°C. Sf9 cells from a 750 mL culture were collected and lysed by resuspending them in 30 mL lysis buffer, followed by incubation on ice for 10 min. A soluble fraction was obtained after centrifugation of the lysate at 15,000×g for 15 min at 4°C. The soluble fraction was incubated with 250 μL anti-FLAG M2 affinity resin equilibrated with lysis buffer for 4 h at 4°C. The beads were collected and washed with 10 mL of wash buffer and then with 5 mL of EB (20 mM HEPES-KOH [pH 7.5], 100 mM KCl, 5 mM MgCl_2_) containing 1 mM DTT. Each recombinant protein was eluted twice in 250 μL EB containing 1 mM DTT and 250 μg/mL 3×FLAG peptide (Sigma-Aldrich). Eluates were pooled and concentrated using a Vivaspin 500 (GE Healthcare).

### Cell culture and cell line generation

Dppa3_KO/UHRF1-GFP mESCs were described previously(35, 51) and were cultivated in Dulbecco’s Modified Eagle’s Medium-High Glucose supplemented with homemade recombinant LIF(52), 16% Fetal Bovine Serum (FBS, Sigma) 0.1 mM β-mercaptoethanol ready to use, (Life Technologies), 2 mM L-glutamine, respectively 100 U/mL and 100 μg/mL of Pen/Strep (Sigma), 1×non-essential amino acids (Sigma) and with 2i inhibitors (1 μM PD32591 and 3 μM CHIR99021, Axon Medchem). Cells were cultured in gelatine (0.2%) coated Corning dishes at 37°C in a 5% CO_2_ incubator and gently dissociated for passaging at 1:8 ratio every second day with Stem Pro Accutase (Gibco). Cells were washed with Dulbecco’s Phosphate Buffer Saline cell culture grade.

To generate stable doxycycline inducible Dppa3 mESC lines, 400,000 Dppa3_KO/UHRF1-GFP mESCs were seeded into one well of a 6-well plate 4 h before transfection. Then cells were transfected with equimolar amounts of pSBtet-3×FLAG- Dppa3_wt-mScarlet-PuroR or pSBtet-3×FLAG-Dppa3_mutants-mScarlet-PuroR or pSBtet- 3×FLAG-B.Taurus Dppa3-mScarlet-PuroR, and the Sleeping Beauty transposase, pCMV(CAT)T7-SB100(53) (Addgene plasmid #34879) vector using Lipofectamine 3000 (Thermo Fisher Scientific) according to manufacturer’s instructions. After 48 h transfection, cells were seeded onto p100 plates into clonal density under puromycin selection with a concentration of 1 μg/mL for 7 days. After the seventh day selection, colonies with mScarlet signal under induction of doxycycline (1 μg/mL) were picked and mixed for further analyses. The stable cell lines were kept in puromycin selection of 1 μg/mL starting two weeks after expansion.

### Sleeping beauty constructs

To generate inducible mouse Dppa3 mutants and Bos Taurus Dppa3 expression constructs, the sequences coding for mDppa3 mutants, including R104A, R89A/V90A, L91A/V94G, M102P/E109P, and Bos Taurus Dppa3 were synthesized as gBlocks (IDT, Coralville, IA, USA) and inserted into the pSB_Avi_3×FLAG_insert_mScarlet_Puro vector (linearized by AsiSI and NotI) using Gibson assembly based on protocol instructions of 2×HiFi master mix (New England Biolabs). The correct constructs were selected based on Sanger Sequencing results (Eurofins).

### Cellular fractionation and western blotting

Cell fractionation was performed as described previously with minor modifications(54). Approximately 1×10^7^ mESCs were resuspended in 100 μL of buffer A (10 mM HEPES [pH 7.9], 10 mM KCl, 1.5 mM MgCl_2_, 340 mM sucrose, 10% glycerol, 0.1% Triton X-100, 1 mM DTT, 1 mM phenylmethylsulfonyl fluoride (PMSF), 1×mammalian protease inhibitor cocktail (PI; Roche)) and incubated for 5 min on ice. Nuclei were collected by centrifugation (5 min, 1300×g, 4°C) and the cytoplasmic fraction (supernatant) was cleared again by centrifugation (15 min, 20,000×g, 4°C). Nuclei were washed once with buffer A and resuspended in 100 μL of buffer A. The cytoplasmic and nuclear fractions were supplemented with 4×Laemmli buffer and boiled for10 min at 95°C.

Western blots were performed using the following antibodies: anti-alpha-Tubulin (monoclonal; 1:2000; Sigma, T9026), rabbit anti-H3 (polyclonal; 1:5000; Abcam, ab1791), mouse anti-GFP (monoclonal; 1:5000; Roche), goat anti-rabbit HRP (polyclonal; 1:5000; BioRad), rabbit anti-mouse HRP (1:5000; Invitrogen, Cat # A27025).

### Live cell imaging

Live cell experiments were conducted with a Nikon TiE microscope equipped with a Yokogawa CSU-W1 spinning disk confocal unit (pinhole size 50 μm), together with a Andor Borealis illumination unit, [Andor ALC600 laser combiner (405 nm/ 488 nm/ 561 nm/ 640 nm)]. The images were acquired with an Andor IXON 888 Ultra EMCCD camera, with 100×/1,45NA oil immersion objective through the interface of the software NIS Elements (Version 5.02.00) in Perfect Focus System with lasers at 488 nm for GFP, 561 nm for mScarlet and 640 nm for Sir-DNA. For live cell imaging, 80,000 cells were seeded into one well of an 8 well glass bottom chamber slide (Ibidi) coated one day before with Geltrex Ready-to-Use (Thermo Fisher Scientific). The next day, 1 μM Sir-DNA (Spirochrome) for DNA staining was added into medium 1 h before live cell imaging. Images were acquired before and after 50 min of doxycycline induction with the same laser power, acquisition time and gain.

### Targeted bisulfite amplicon sequencing (TaBAseq)

TaBAseq was performed as described previously(35). According to the manufacturer’s instructions, there were 1×106 ESCs culturing in 2iLIF condition for genomic isolation with QIAamp DNA Mini Kit (Qiagen). 500 ng of gDNA was used for bisulfite conversion followed by the instructions of EZ DNA Methylation-Gold Kit (Zymo), which was eluted in a 2×20 μL Elution Buffer.

TaBAseq is based on two sequential PCRs. The first one amplifies locus-specific LINE-1 elements and the second one indexes the sample-specific amplicon with Ilumina’s Truseq and Nextera compatible overhangs. The amplicon-specific PCR was performed in a total volume of 25 μL containing 0.4 μM of forward and reverse primers, 2 mM betaiinitialne (Sigma-Aldrich), 10mM tetramethylammonium chloride solution (Sigma-Aldrich), 1×MyTaq Reaction Buffer, 0.5 units of MyTaq HS polymerase (Bioline) and 1 μL of bisulfite converted DNA (12.5 ng). The cycling parameters were as follows: 5 min at 95°C for initial denaturation, 40 cycles (95°C for 20 s, 58°C for 30 s, 72°C for 25 s) and with the final elongation 72°C for 3 min. To check for the quality and yield of the PCR reaction, it was run on a 2% agarose gel and to purify the PCR amplicons, it was used CleanPCR beads with 1.8 × the volume of the remaining PCR reaction. The magnetic beads were immobilized with DynaMag-96 Side Magnet (Thermo Fisher) for 5 min. After the supernatant is removed, the beads are washed 2 × with 150 μL of fresh 70 % ethanol. After leaving the beads to air-dry for 5 min, DNA was eluted in 15 μL of elution buffer (10 mM Tris-HCl [pH 8.0]). DNA concentration was determined using Nanodrop (Thermo Fisher Scientific).

To index the amplicons, a second PCR was performed in a 25 μL total volume, containing 0.08 μM (1 μL of a 2 μM stock) of the Indexing Primers, respectively i5 and i7, 1×MyTaq Reaction Buffer, 0.5 units of MyTaq HS Polymerase (Bioline) and 1 μL of the purified PCR purified amplicon. The cycling parameters are as follows: 5 min at 95°C for initial denaturation, 18 cycles (95°C for 20 s, 55°C for 30 s, 72°C for 40 s) and with the final elongation at 72°C for 5 min. Again 2% agarose gel was used to determine the yield and purity of the PCRs. The PCR-indexed amplicons were purified as described above using the CleanPCR magnetic beads. The DNA concentration for each sample was determined with Quant-iT PicoGreen dsDNA Assay Kit (Thermo Fisher Scientific). The final library was created by pooling together an equimolar ratio of all PCR products in a final concentration of 1 ng/μL of the total library. The final library concentration was once again estimated via Quant-iT PicoGreen, while the size distribution and quality of the library were assessed using Bioanalyzer. Ilumina Miseq was used to sequence the dual-indexed TaBAseq library with a 2×300 bp paired-end, with 5% sequencing coverage. The sequencing data was analyzed with a TABSAT package(55).

### Protein sequence alignments

Protein sequence alignments for UHRF1 and DPPA3 were performed using Jalview software (www.jalview.org) with sequences retrieved from Uniprot for UHRF1: (Q8VDF2_Mus musculus; Q7TPK1_Rattus norvegicus; G3RVG5_Gorilla gorilla gorilla; Q96T88_Homo sapiens; H2QF26_Pan troglodytes; A0A3Q7TCY9_Vulpes vulpes; A0A3Q7V1Z9_Ursus arctos horribilis; A0A2U3VKN8_Odobenus rosmarus divergence; A0A2Y9KPL3_Enhydra lutris kenyoni; A7E320_Bos Taurus) and DPPA3 (Q8QZY3_Mus musculus; Q6IMK0_Rattus norvegicus; G3RB81_Gorilla gorilla gorilla; Q6W0C5_Homo sapiens; H2Q5C8_Pan troglodytes; A0A3Q7U513_Vulpes vulpes; A0A3Q7W0Q7_Ursus arctos horribilis; A0A2U3ZR98_Odobenus rosmarus divergence; A0A2Y9IX83_Enhydra lutris kenyoni; A9Q1J7_ Bos Taurus). Sequences are aligned with the multiple sequence alignment algorithm MAFFT (http://mafft.cbrc.jp/alignment/software/). The percentage of protein identity conserved for both proteins is calculated through Pairwise alignment algorithm between Mus musculus and different species. The phylogenetic tree was generated with online tools, https://www.genome.jp (PhyML) and iTOL.

## RESULTS

### Biochemical assay for determining essential regions for complex formation

The C-terminal fragment of mDPPA3 is required for binding to mUHRF1 (Figure 1A, B)(36). GST pull-down assays using truncated mouse UHRF1 showed that the PHD finger (residues 304–372: mPHD) was sufficient for binding to the C-terminal fragment of mDPPA3 (residues 61–150) (Supplementary Figure S1A, B). Further optimization revealed that mDPPA3 residues 76–127, followed by an additional Trp (mDPPA3_76–128W_), were sufficient to bind to the mPHD (Figure 1C). This interaction was validated by isothermal titration calorimetry (ITC) and NMR spectroscopy. mDPPA3_76–128W_ bound to mPHD with a *K*_D_ of 27.7 nM, which is stronger than the binding of the N-terminal tail of H3 (*K*_D_: 1590 nM) and PAF15 (*K*_D_: 3523 nM) to mPHD, which are well-known ligands of the UHRF1 PHD finger (Figure 1D, Supplementary Figure S2). The ^1^H-^15^N heteronuclear single quantum coherence (HSQC) spectrum of [^15^N]-mPHD titrated with non-labeled mDPPA3_76–128W_ was markedly different from that of apo-mPHD, in which nearly all HSQC signals were shifted in the slow-exchange regime on the chemical shift timescale (Supplementary Figure S3A, B). The weighted averages of the ^1^H and ^15^N chemical shift differences (Δδ) between the apo-mPHD and mPHD bound to mDPPA3 showed relatively large values for Glu333, Leu337, Met343, and Glu362 in mPHD (Supplementary Figure S3A–C). ITC data indicated that the L337A mutation in mPHD severely weakened the binding to mDPPA3_76–128W_ with a *K*_D_ > 43,000 nM (Figure 1D, Supplementary Figure S2) and also abolished the binding to the N-terminal tail of H3 and PAF15 (Supplementary Figure S2). The ^1^H-^15^N HSQC spectrum of mPHD L337A showed that the mutation had a modest effect on the native conformation (Supplementary Figure S3D). Collectively, Leu337 of mPHD is commonly used as the interface for binding to mDPPA3, H3, and PAF15.

### Overall structure of mPHD in complex with mDPPA3

To uncover the molecular mechanism by which mDPPA3 binds to mPHD with high affinity, we determined the structure of the complex using solution NMR analysis. As a target for structural analysis, we designed a chimeric protein, mDPPA3_76–127_ was fused to the C-terminal end of mPHD (mPHD-mDPPA3). The ^1^H-^15^N HSQC spectrum of the mPHD moiety in [^15^N]-mPHD-mDPPA3 was superimposed on that of [^15^N]-mPHD mixed with non-labeled mDPPA3_76–128W_, validating that the chimeric protein has a binding mode equal to that of the isolated proteins (Supplementary Figure S4A).

The ensemble of the mPHD-mDPPA3 structures was well converged and showed a low average root mean square deviation (rmsd) of 0.40 Å for Cα atom coordinates (Figure 2A, B, Table 1 and 2). The mPHD moiety comprised of pre-PHD (residues 304–321) and core-PHD (residues 322–372) encompassing three zinc finger motifs (Zn1-3) and an anti-parallel β-sheet (Figure 2A). The residues 307–369 of the mPHD moiety were superimposed on the solution structure of apo-mPHD (PDB: 6VFO) with a Cα rmsd of 3.1 Å (Supplementary Figure S5), indicating that the overall structure of mPHD did not change substantially upon binding of mDPPA3, except for a loop region, as mentioned later.

**Figure 2:**
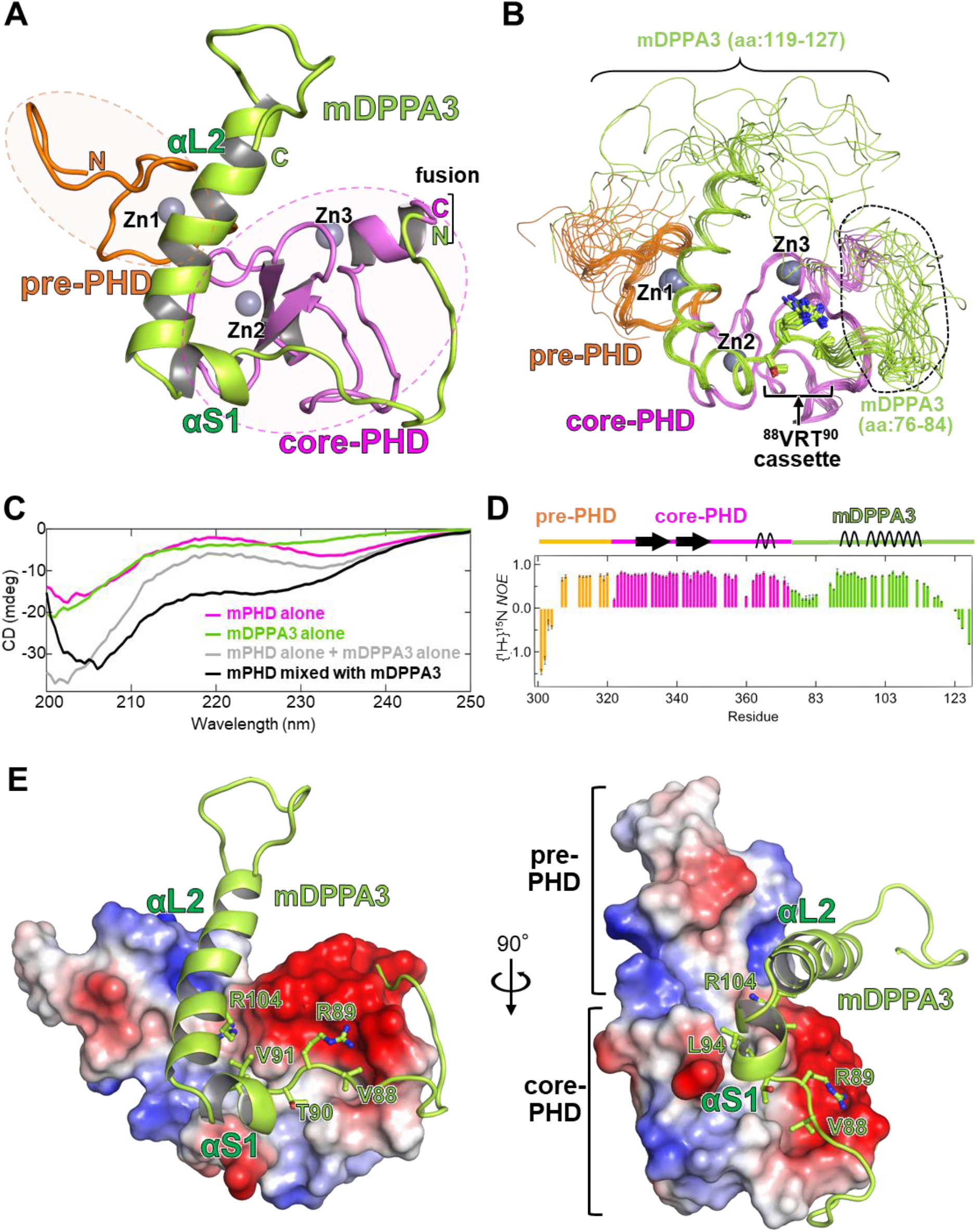
Structural analysis of mPHD-mDPPA3. **(A)** Overall NMR structure of mPHD-mDPPA3. Pre-PHD, core-PHD, and mDPPA3 are shown as orange, pink, and green cartoons, respectively. Zinc ions are depicted as gray-sphere models. **(B)** Overlay of 20 NMR structures in which the structurally unconverged regions of mDPPA3 are indicated. The color scheme is the same as that in Figure 2A. **(C)** CD spectra of mPHD alone (pink), mDPPA3_76–128W_ alone (green), and the mPHD in complex with mDPPA3_76–128W_ (black). The sum of CD spectra of mPHD alone and mPDDA3_76–128W_ alone is shown as gray. **(D)** Backbone {^1^H}-^15^N heteronuclear NOE of mPHD-mDPPA3. The het-NOE values were color-coded orange and pink for mPHD and green for mDPPA3_76–127_ regions. **(E)** Electrostatic surface potential of mPHD. The surface colors red and blue represent negative and positive charges, respectively. mDPPA3 is depicted as a green cartoon with a stick model of the key residues that interact with mPHD.

The ^1^H-^15^N HSQC spectrum of [^15^N]-mDPPA3 in the free state showed sharp signals in a very narrow range of ^1^H chemical shifts corresponding to a randomly coiled state, indicating the unstructured conformation of apo-mDPPA3 (Supplementary Figure S4B). Binding of mPHD induced two α-helices in mDPPA3: a short α-helix, αS1 (residues 90–95) and a long α-helix αL2 (residues 97–118), which are connected via a short turn, resulting in an ‘L’ shaped motif in mDPPA3 (Figure 2A, B). The helical structure induced upon binding of mPHD was further experimentally confirmed by NMR and circular dichroism (CD) spectrum analysis. Secondary structure prediction by TALOS-N using the backbone ^1^HN, ^15^N, ^13^C_α_, ^13^C_β_, and ^13^CO chemical shifts showed the induction of helical structure of mDPPA3 upon binding of mPHD (Supplementary Figure S4C, D)(56). The CD spectrum of mPHD mixed with mDPPA3 showed a negative peak at 222 nm, whereas this property was not observed in the spectra of mPHD and mDPPA3 alone (Figure 2C). ^1^H-^15^N heteronuclear nuclear Overhauser effect (het-NOE) of [^15^N]-mPHD-mDPPA3, which is sensitive to local conformational flexibility on the picosecond to nanosecond timescale(57), was consistently high except for the residues on the N-terminal region of the mPHD moiety, residues 76–84, and the C-terminus of mDPPA3 (Figure 2B, D). The NMR structure showed that residues 85–118 of mDPPA3 in the complex were structurally well converged (Figure 2B) and ITC data demonstrated that mDPPA3_85–118_ bound to mPHD with a *K*_D_ = 45 nM, which is comparable to the binding affinity of mDPPA3_76–128W_, indicating that residues 85–118 of mDPPA3 were sufficient for binding to mPHD (Supplementary Figure S2). mDPPA3 provides a wide interface for contact with mPHD; the estimated contact area between the two proteins was ∼1360 Å^2^, which comprises three parts: a ^88^VRT^90^ cassette, and αS1 and αL2 of mDPPA3 (Figure 2E). In contrast, H3 and PAF15 bind to the human UHRF1 PHD finger with a contact area of ∼400 Å^2^ (PDB: 3ASL) and ∼360 Å^2^ (PDB: 6IIW), respectively. The contact areas of the complex structures are much smaller than those of the mPHD:mDPPA3 complex, supporting the stronger binding affinity between mPHD and mDPPA3. NMR titration experiments demonstrated that mDPPA3 preferentially bound to mPHD even in the presence of excess H3 peptide (Supplementary Figure S4E), indicating that mDPPA3-binding to mPHD competes with the H3-binding.

### Recognition of the conserved VRT cassette in mDPPA3 by mPHD

The ^88^VRT^90^ cassette of mDPPA3 adopted an extended conformation and fit into the shallow acidic groove on mPHD via an intermolecular anti-parallel β sheet (Figure 2A). The side chain of Val88 in the ^88^VRT^90^ cassette was surrounded by Leu336, Val357, Pro358, Glu360, and Trp363 of mPHD within van der Waals contacts (Figure 3A). The guanidino group of Arg89 in the cassette was located within a distance that enabled hydrogen bonding with the side-chain carboxyl groups of Asp339 and Glu362 of mPHD (Figure 3A). The side-chain hydroxyl and methyl groups of Thr90 in mDPPA3 interacted with mPHD differently: the methyl group formed hydrophobic interactions with Leu336, Leu337, and Val357 of mPHD, whereas the hydroxyl group was positioned within the hydrogen bond distance with the main chain amide of Ser93, indicating its function as a helix-cap for the N-terminus of the following αS1 (Figure 3B). ITC data showed that the R89A/T90A mutations in the ^88^VRT^90^ cassette of mDPPA3 abolished its interaction with mPHD (Figure 3C). This interaction is supported by GST pull-down experiments with *in vitro* translated full-length mUHRF1 and GST-mDPPA3, which showed that the R89A/T90A mutations are sufficient to block the binding of mDPPA3 to mUHRF1 (Figure 3D), indicating that the ^88^VRT^90^ cassette in mDPPA3 plays a critical role in binding to mPHD.

**Figure 3:**
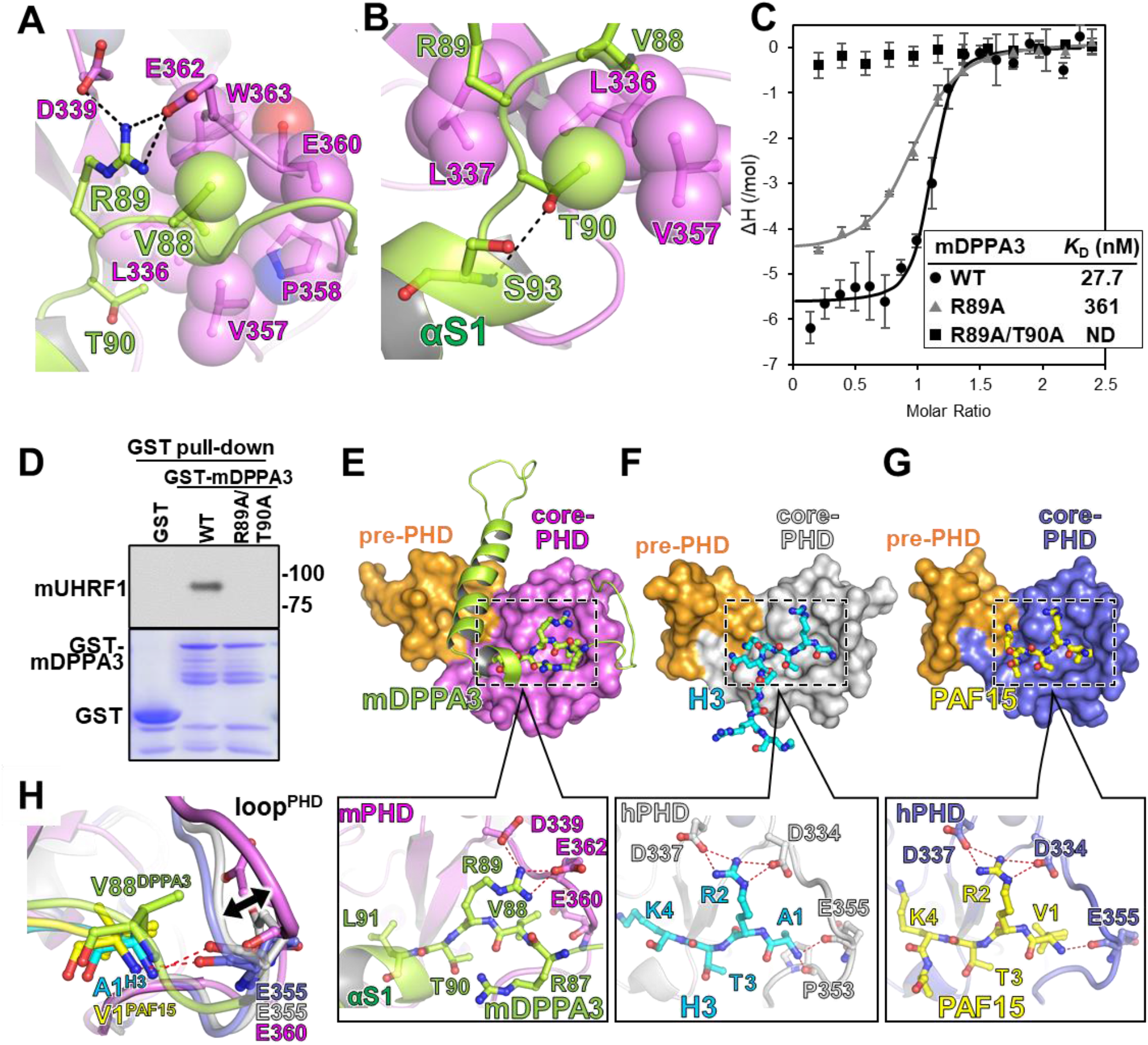
Recognition of the ^88^VRT^90^ cassette of mDPPA3 by mPHD. **(A)** Recognition of Val88 and Arg89 in mDPPA3. The mDPPA3 residues are shown as a green stick model and transparent sphere model of Val88 methyl groups overlaid on the stick model. mPHD residues that are involved in the recognition of Val88 of mDPPA3 are depicted as a pink stick model superimposed on a transparent sphere model, and Asp339/Glu362 are shown as pink stick models. The black dashed lines indicate the hydrogen bonds. **(B)** Recognition of Thr90 of mDPPA3 showing a green stick model for hydrophobic residues of mPHD and a pink stick model with a transparent sphere model. The hydrogen bond is indicated by the black dashed line. **(C)** ITC measurements using mutants in the VRT cassette of mDPPA3 and mPHD WT. Superimposition of enthalpy change plots with standard deviations. **(D)** GST pull-down assay to detect the interaction between full-length mUHRF1 and full-length GST-mDPPA3 wild-type (WT) and mutant proteins. **(E)** The upper panel shows the overall structure of mPHD (pre-PHD: orange surface, core-PHD: pink surface) bound to mDPPA3 (green stick model). The lower panel shows recognition of the ^88^VRT^90^ cassette of mDPPA3. The red dashed lines indicate the hydrogen bonds. **(F)** The upper panel shows the overall structure of hPHD (pre-PHD: orange surface, core-PHD: gray surface) bound to H3 (cyan stick model) (PDB: 3ASL). The lower panel shows recognition of ^1^ARTK^4^ of H3 by human PHD. **(G)** The upper panel shows the overall structure of hPHD (pre-PHD: orange surface, core-PHD: light-purple surface) bound to PAF15 (yellow stick model) (PDB: 6IIW). The lower panel shows recognition of ^1^VRTK^4^ of PAF15 by hPHD. **(H)** Overlay of the N-terminus of H3, PAF15, and V88 of mDPPA3. The double arrow indicates the structural difference between the loops in the PHDs.

The ^88^VRT^90^ cassette of mDPPA3 was well-conserved in the N-terminal sequences of PAF15 (^1^VRT^3^) and H3 (^1^ART^3^). Although the ^88^VRT^90^ cassette of mDPPA3 is not located at its N-terminal end, the cassette was recognized by the shallow acidic groove of mPHD, which is used for PAF15/H3-binding (Figure 3E–G). Notably, compared with the PHD moiety structure in the complexes with PAF15/H3 and mDPPA3, the conformation of loop residues 356–364 (hereafter loop^PHD^) in mPHD was markedly different (Figure 3H). In the complex with PAF15/H3, loop^PHD^ functions as a wall to recognize the N-terminal amino group of PAF15/H3 by hydrogen bonding using the main chain carbonyl oxygen of Glu355 (Figure 3F, G). In contrast, in the complex with mDPPA3, loop^PHD^ is shifted outward away from the amide nitrogen of Val88 of mDPPA3, resulting in disruption of the hydrogen bond between the amide nitrogen in mDPPA3 and the main-chain carbonyl oxygen of Glu360 (corresponding to residue Glu355 in humans) in the loop^PHD^ of mPHD (Figure 3H). The results indicate that loop^PHD^ has an intrinsically flexible capability to accommodate the VRT cassette of mDPPA3. The structural rearrangement of loop^PHD^ permits the peptide bond moiety between Arg87–Val88 to enter the groove (Figure 3E, H). The side-chain conformation of Arg89 of mDPPA3 in the complex with mPHD was different from that of Arg2 of H3/PAF15 (Figure 3E–G). Given that ITC data for the R89A mutation of mDPPA3 showed an ∼18-fold decrease in binding affinity to mPHD (*K*_D_ = 361 nM; Figure 3C, Supplementary Figure S2), the Arg89 guanidino group of mDPPA3 interacts with the acidic surface of mPHD.

### Binding mode of two-helices in mDPPA3 to mPHD

In addition to the ^88^VRT^90^ cassette, mDPPA3 utilizes two α-helices, αS1 and αL2, to bind to mPHD. The αS1 of mDPPA3 forms a small hydrophobic cluster composed of the side chain methyl groups of Leu91, Val94, and Leu95, of which Leu91 and Val94 interact with the hydrophobic patch of mPHD comprising Leu337 and Ala344 (Figure 4A). As mentioned before, the L337A mutation in mPHD largely impaired the interaction with mDPPA3; similarly, the L91A/V94A mutations in mDPPA3 also reduced the binding affinity to mPHD with a *K*_D_ of 1084 nM (Figure 4B), indicating that αS1 contributes to stable complex formation.

**Figure 4:**
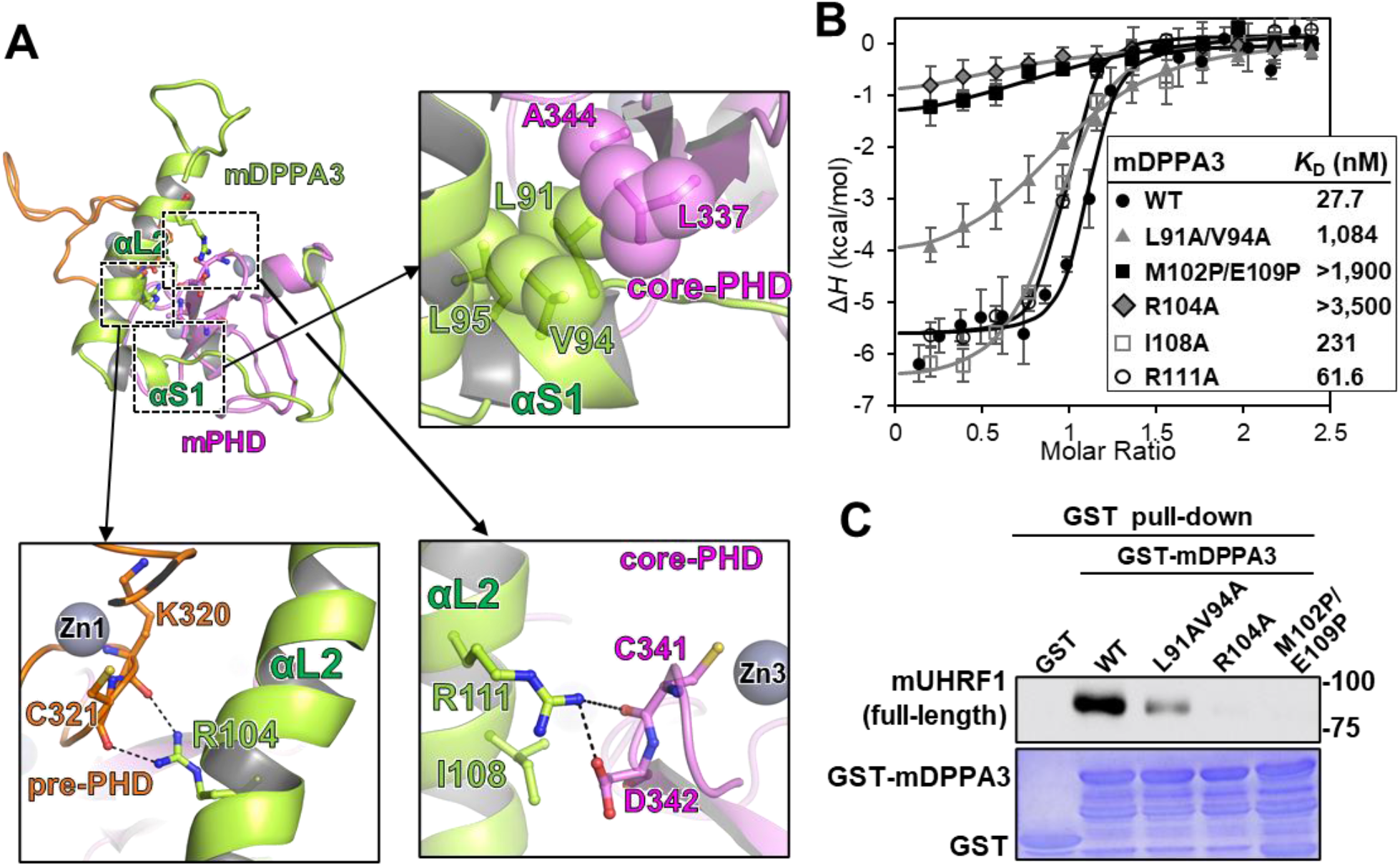
Contribution of the two helices of mDPPA3 for binding to mPHD. **(A)** Interaction between the two helices of mDPPA3 and mPHD. The top-left panel displays the overall structure of the mPHD-mDPPA3 complex in the cartoon mode. The top-right panel shows the interaction between the αS1 of mDPPA3 and the hydrophobic patch of mPHD. The bottom-left panel depicts the binding of Arg104 in mDPPA3 to the pre-PHD domain. The black dashed lines indicate the hydrogen bonds. The bottom-right panel shows the interaction between Cys341/Asp342 in core-PHD and Ile108/Arg111 on the αL2 of mDPPA3. The color scheme is the same as that shown in Figure 3A. Transparent sphere models of the side chains of hydrophobic residues were superimposed on the corresponding stick models. **(B)** ITC measurements using mutants of α-helices of mDPPA3 and WT mPHD. Superimposition of enthalpy change plots with standard deviations. **(C)** GST pull-down assay to detect the interaction between full-length mUHRF1 and GST-mDPPA3 WT and mutants in αS1 and αL2.

αL2 of mDPPA3, comprising residues 99–118, was embedded into a concave surface between the pre- and core-PHD domains (Figure 2E). Notably, the introduction of the helix breaking residue M102P/E109P mutations, or truncation of αL2 residues 76–106 in mDPPA3 resulted in a significant reduction in binding to mPHD, indicating that the formation of αL2 is critical for interaction with mPHD (Figure 4B, Supplementary Figure S2). Interestingly, the Arg104 guanidino group on αL2 in mDPPA3 was deeply buried in the binding interface of the complex, in which the side chain formed hydrogen bonds with the main chain carbonyl oxygens of Lys320 and Cys321 in the pre-PHD domain (Figure 4A). Ile108 and Arg111 of mDPPA3 also interacted with Cys341 and Asp342 of the core-PHD domain, indicating that αL2 bridges the pre- and core-PHD domains (Figure 4A). Although mutations of Ile108 and Arg111 of mDPPA3 had a limited effect on binding to mPHD (Figure 4B, Supplementary Figure S2), the R104A mutation significantly reduced the binding affinity to mPHD (Figure 4B). GST pull-down experiments demonstrated that the L91A/V94A mutation in αS1 reduced the binding of GST-mDPPA3 to full-length mUHRF1 modestly (Figure 4C), showing a limited effect on full-length proteins. In contrast, the M102P/E109P and R104A mutations in αL2 of mDPPA3 markedly reduced binding to mUHRF1 (Figure 4C), indicating that αL2 of mDPPA3 plays a pivotal role in complex formation with full-length proteins.

### Mutational analyses of interaction interface of DPPA3 with UHRF1 in *Xenopus* egg extracts and mESCs

To test the function of the DPPA3 mutants, we measured their ability to inhibit UHRF1-dependent maintenance of DNA methylation in *Xenopus* egg extracts (Figure 5A). We compared the inhibitory activity of GST-mDPPA3 (wild-type) with that of the R89A/T90A, L91A/V94A, M102P/E109P, and R104A mutants. Our results showed that wild-type mDPPA3 inhibited the chromatin recruitment of UHRF1 and DNA methylation; in contrast, the R89A/T90A mutant showed a severely impaired inhibitory effect (Figure 5B, C). Concordant with the ITC data and GST-pull down assay, the L91A/V94A mutant was ineffective in inhibiting UHRF1 (Figure 5B). Mutants linked to the αL2 helix structure (M102P/E109P and R104A) were unable to inhibit UHRF1 chromatin loading and DNA methylation (Figure 5B, C). In addition, *in vitro* ubiquitination assay using C-terminal FLAG tagged H3 1-37W as a substrate demonstrated that WT mDPPA3 inhibited the ubiquitination of H3 catalyzed by UHRF1, however, the mutants mDPPA3 failed to do so (Supplementary Figure S6), suggesting that DPPA3 represses the E3-ligase activity of UHRF1.

**Figure 5:**
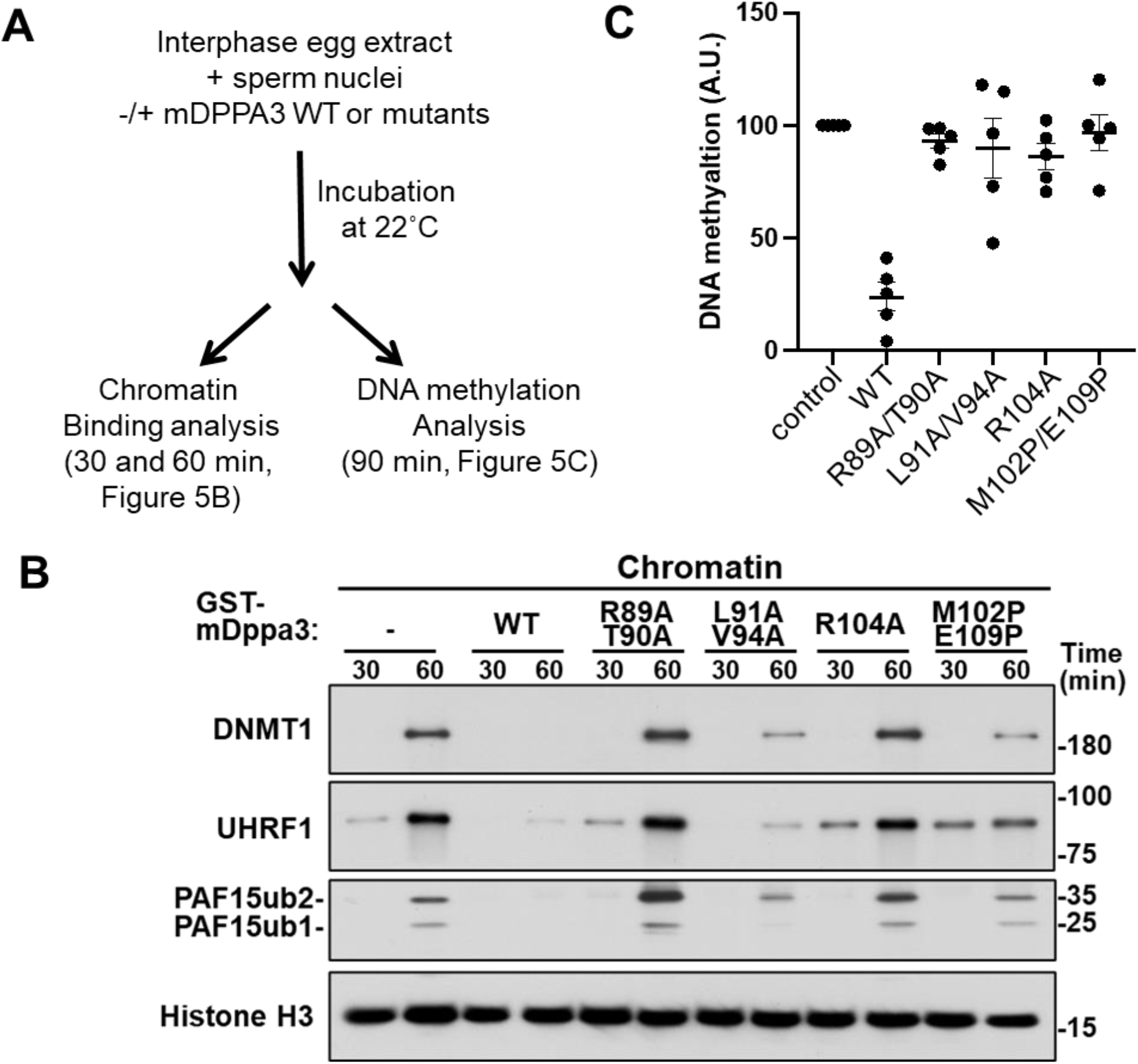
The interaction interface between mDPPA3 and UHRF1 is important for the inhibition of DNA methylation and UHRF1 chromatin binding in *Xenopus* egg extracts. **(A)** Experimental design for functional analysis of mDPPA3 mutants using *Xenopus* egg extracts. **(B)** Sperm chromatin was incubated with interphase *Xenopus* egg extracts supplemented with buffer (+buffer), GST-mDPPA3 (+mDPPA3), or each mDPPA3 mutant. Chromatin fractions were isolated and immunoblotted using the indicated antibodies. Representative data from n = 3 independent experiments. **(C)** Sperm chromatin was added to interphase egg extracts supplemented with radiolabeled S-[methyl-^3^H]-adenosyl-L-methionine and WT-GST-mDPPA3 or each mDPPA3 mutants. The efficiency of maintenance DNA methylation was assessed by the incorporation of radio-labeled methyl groups from S-[methyl-^3^H]-adenosyl-L-methionine (^3^H-SAM) into DNA purified from egg extracts.

To further examine the importance of the interaction interface of DPPA3 and UHRF1, we analyzed chromatin displacement and nucleocytoplasmic translocation of UHRF1 by inducing the expression of DPPA3 mutants in mouse embryonic stem cells (mESCs). We generated inducible mDPPA3-mScarlet expression cassettes harboring mutations R89A/T90A, L91A/V94G, R104A, and M102P/E109P. After introducing these expression cassettes into DPPA3 knock out/UHRF1-GFP (D3KO/U1GFP) mESCs, we used live-cell imaging to observe the localization of UHRF1-GFP (Figure 6A). Wild-type mDPPA3 caused chromatin displacement and nucleocytoplasmic translocation of UHRF1-GFP. In contrast, the mDPPA3 mutants failed to efficiently re-localize UHRF1-GFP (Figure 6A). Furthermore, biochemical fractionation experiments showed that UHRF1-GFP was detected in the cytoplasm after induction of wild-type mDPPA3, whereas the mDPPA3 mutants showed low activity for UHRF1-GFP displacement and export to the cytoplasm. (Figure 6B). The L91A/V94G(A) mutation of mDPPA3 had a more severe effect on mESCs than on *Xenopus* egg extracts (Figure 5B and 6B). We examined the ability of wild-type DPPA3 and the mutants to rescue the hypermethylation of LINE-1 (long interspersed nuclear element) elements in DPPA3 knock-out mESCs (Figure 6C). The mDPPA3 mutants failed to restore low methylation levels (although the M102P/E109P mutant was somewhat less effective) (Figure 6C), indicating that binding of mDPPA3 to UHRF1 PHD finger is required for DNA demethylation.

**Figure 6:**
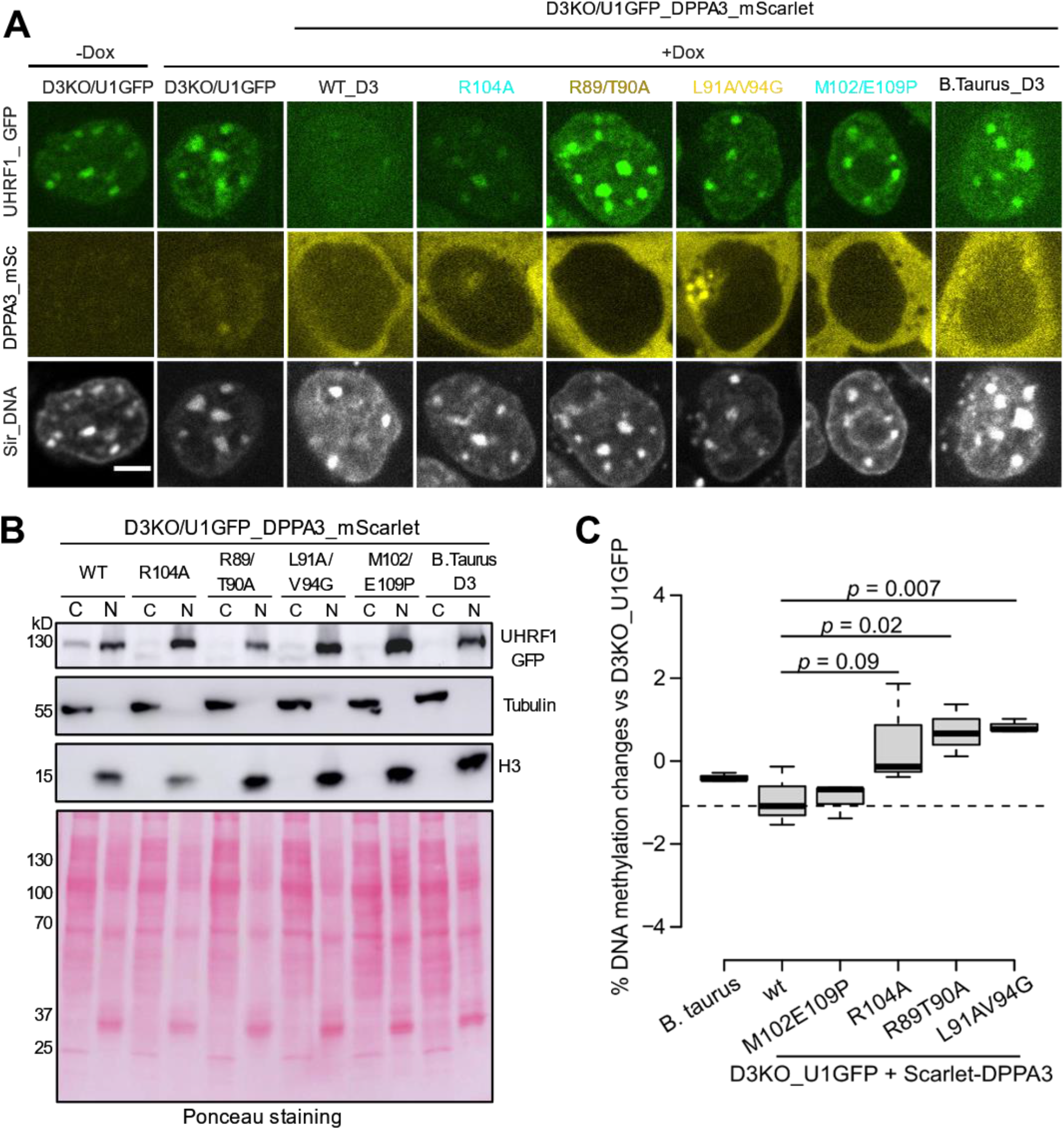
Effect of nuclear localization of mUHRF1 by mDPPA3 mutants in mouse ESCs. **(A)** Representative images illustrating the localization of UHRF1-GFP, and mouse and *B. taurus* DPPA3-mScarlet fusions in live U1G/D3KO + pSB-D3-mSC ESCs after doxycycline induction. DNA counterstain: SiR-DNA. Scale bar: 5 μm. **(B)** Subcellular distribution of UHRF1-GFP before and after DPPA3-mScarlet induction as determined using cell fractionation and western blot analyses. Cells were fractionated into cytoplasmic (C) and nuclear fractions (N). Anti-tubulin and anti-H3 blots were performed to identify the two fractions. Ponceau S stained blots were used as a loading control. UHRF1-GFP was detected using an anti-GFP antibody to determine its distribution in the C and N fractions. **(C)** DNA methylation of LINE-1 repetitive elements were measured by targeted amplicon bisulfite sequencing (TaBAseq) for each cell line (n=3). In the boxplot, the horizontal lines represent the median values, box limits indicate the 25th and 75th percentiles as determined by R software; whiskers extend to minimum and maximum values. A one-tailed t-test was performed and p values are indicated.

Next, we examined the evolutionary conservation of the DPPA3-UHRF1 interaction interface. Although UHRF1 is highly conserved, phylogenetic analysis of DPPA3 showed the prevalence of two divergent groups: group 1, with more than 24.6% similarity (*Mus musculus, Rattus norvegicus, Gorilla gorilla, Homo sapiens*, and *Pan troglodytes*) and group 2 which showed less than 18.6% identity (*Vulpes vulpes, Ursus arctos horribilis, Odobenus rosmarus, Enhydra lutris kenyoni*, and *Bos taurus*) (Supplementary Figure S7), suggesting rapid evolution of DPPA3. This is consistent with the observation that DPPA3 has recently evolved in mammals and does not appear to have catalytic activity(35). Notably, the VRT cassette of DPPA3, a central element in the interaction with UHRF1, was highly conserved except in *Bos taurus* (Figure 1B), which could not displace UHRF1 in mESCs from chromatin (Figure 6A).

Collectively, the results of the functional assays indicate that the key residues of mDPPA3 identified in our NMR structure of the complex are important for UHRF1 regulation.

## DISCUSSION

The PHD finger is a well-known reader domain for post-translational modifications of histone H3(58). Several structural studies of PHD fingers in complex with their ligands have revealed an N-terminal recognition rule for ligand recognition that can be applied to the vast majority of PHD fingers. PHD fingers have a shallow acidic groove for recognition of the N-terminus of ligands, in which the amino group of the first amino acid residue in the ligand forms hydrogen bond (s) with the PHD finger, resulting in a binding affinity with *K*_D_ in the μM range. Our NMR solution structure showed that mDPPA3 binds to mPHD via multifaceted interactions, utilizing the VRT cassette and two α-helices. The ^88^VRT^90^ cassette of mDPPA3 is not located at the N-terminal end. However, the cassette was recognized by the shallow acidic groove of mPHD in a similar manner to the N-terminus of ^1^ART^3^ in H3 and ^1^VRT^3^ in PAF15. This binding is reinforced by the two α-helices unique to mDPPA3, resulting in high-affinity binding to mPHD (*K*_D_ = 27.7 nM). Although the N-terminus recognition rule does not apply to mDPPA3 binding to mPHD, stable complex formation is ensured by the multifaceted interaction provided by mDPPA3. The VRT cassette of mDPPA3 is well conserved among various species (Figure 1B), suggesting that binding of the VRT cassette to the UHRF1 PHD finger while disregarding the N-terminus recognition rule is a common molecular mechanism for the complex formation of DPPA3 and UHRF1. Intriguingly, phosphorylation of H3T3 negatively regulates the chromatin binding of UHRF1(59). NetPhos3.1 (https://services.healthtech.dtu.dk/service.php?NetPhos-3.1) predicts the high probability that T90 of mDPPA3 is target for protein kinase A (^86^R<u>R</u>VR<u>T</u>^90^), suggesting that post-translational modifications of mDPPA3 regulate its binding to UHRF1.

The VRT cassette and αS1 helix of DPPA3 identified in the NMR structure were relatively conserved between species, whereas the αL2 helix was less conserved and had a short insertion in the group 2 species of the DPPA3 family (Figure 1B, Supplementary Figure S7). AlphaFold2 predicted that the corresponding regions in humans (UniProt: Q6W0C5) and rats (UniProt: Q6IMK0) show a long helical structure, suggesting that the formation of the αL2 in the DPPA3 family is structurally conserved.

In general, a conventional PHD finger consists of one core-PHD domain, including two zinc fingers(58). In contrast, the UHRF1 PHD finger contains a pre-PHD domain, which provides an additional binding surface for mDPPA3. Protein sequence analysis of UHRF1 comparing 10 different species showed that UHRF1 was highly conserved during mammalian evolution (>40% similarity) (Supplementary Figure S7). In particular, the UHRF1 pre-PHD and PHD domains were very similar (61.1%) (Figure 1B), suggesting the conservation of the domain in the UHRF1 family. The αL2 of mDPPA3 is embedded in the concave surface formed between the pre- and core-PHD domains in mPHD. Breaking the folding of the αL2 helix severely impaired the binding affinity to mPHD, indicating that αL2 formation by mDPPA3 is crucial for its interaction with mPHD. Thus, the unique structural features of mPHD, consisting of pre- and core-PHDs, and of mDPPA3 with the two helices for enforcing VRT cassette binding ensure highly specific structural complementarity for complex formation. As a consequence, DPPA3 does not bind to the conventional PHD finger that lacks the pre-PHD domain.

The binding of the UHRF1 PHD domain to H3 is involved in its chromatin localization(29, 35, 36). The shallow acidic groove on mPHD is a common binding platform for mDPPA3 and H3. In addition to the shallow acidic groove, mDPPA3 utilizes two helices for binding to mPHD, which increases its binding affinity. DPPA3-binding to mPHD totally competes with the H3-binding (Supplementary Figure S4E). Thus, the molecular mechanism by which mDPPA3 inhibits chromatin binding of UHRF1 includes the preferential binding of mDPPA3 to the UHRF1 PHD finger. Indeed, mDPPA3 inhibited the ubiquitination of H3 catalyzed by UHRF1 *in vitro* (Supplementary Figure S6), suggesting that DPPA3 inhibits chromatin binding of UHRF1 and represses the E3-ligase activity of UHRF1 in cells. The binding mode of DPPA3 to UHRF1 is unique and DPPA3-binding explicitly inhibits chromatin loading of UHRF1. UHRF1 is overexpressed in many cancer cells and downregulation of UHRF1 in these cells leads to reactivation of tumor suppressor gene expression(60). Overexpression of DPPA3 leads to tumor differentiation in hepatocellular carcinoma in which UHRF1 nuclear translocation is impeded(61). Thus, our structural analysis of the UHRF1 PHD in complex with DPPA3 may provide a framework for the design of new anticancer drugs. Peptide-like inhibitors that mimic the αL2 of DPPA3 and specifically bind to the concave surface between pre- and core-PHD domains inhibit excessive UHRF1 in cancer cells.

## ACCESSION NUMBERS

NMR data and refined coordinate was deposited in the Protein Data Bank with accession code 7XGA and Biological Magnetic Resonance Bank with accession code bmr36483. Supplementary Information is linked to the online version of the paper at www.nature.com/nature. Correspondence should be addressed to. Source data are provided with this paper.

## Supplementary Data statement

Supplementary Data are available at NAR Online.

## TABLE AND FIGURES LEGENDS

**Supplementary Figure S1:**
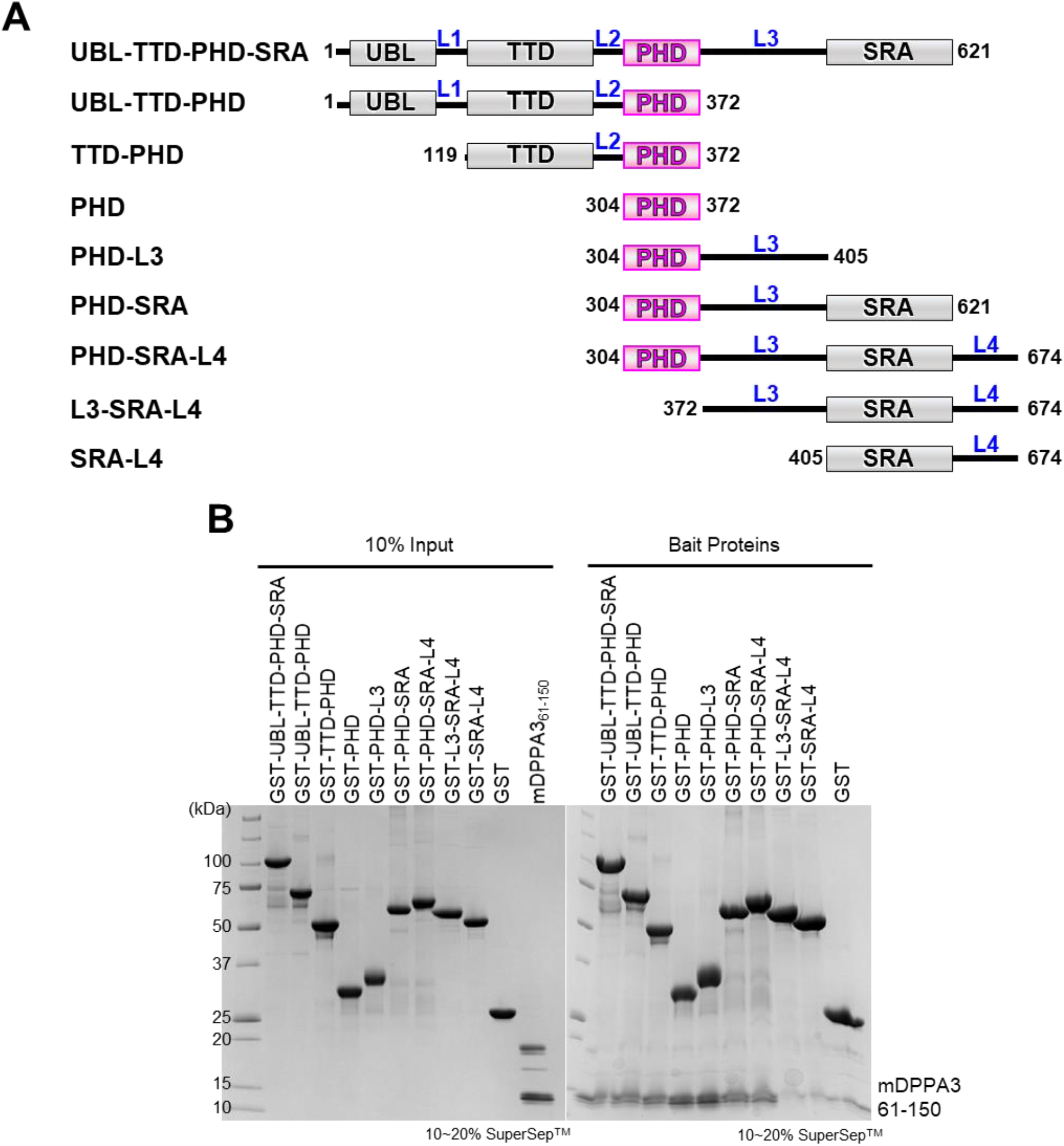
GST pull-down assay for detecting the interaction between truncate version of GST fused mUHRF1 and mDPPA3. **(A)** Schematic figure of domain structures of mUHRF1 that used for the pull-down assay **(B)** GST pull-down experiments using truncated forms of GST-mUHRF1 and C-terminal fragment of mDPPA3 (residues 61–150). Proteins are stained by Coomassie Brilliant Blue.

**Supplementary Figure S2:**
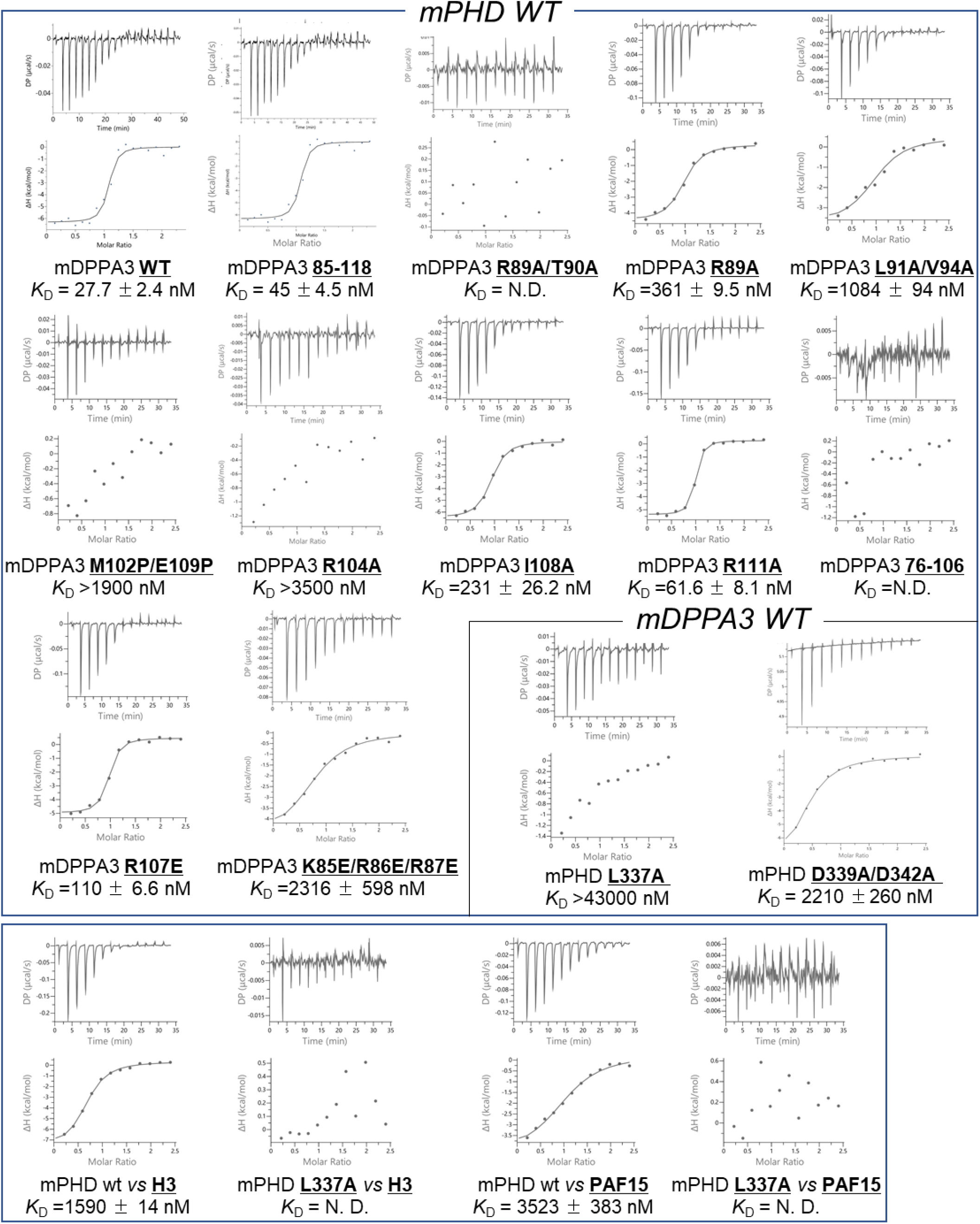
ITC thermograms (upper) and plots of corrected heat values (lower) for the indicated binding experiments.

**Supplementary Figure S3:**
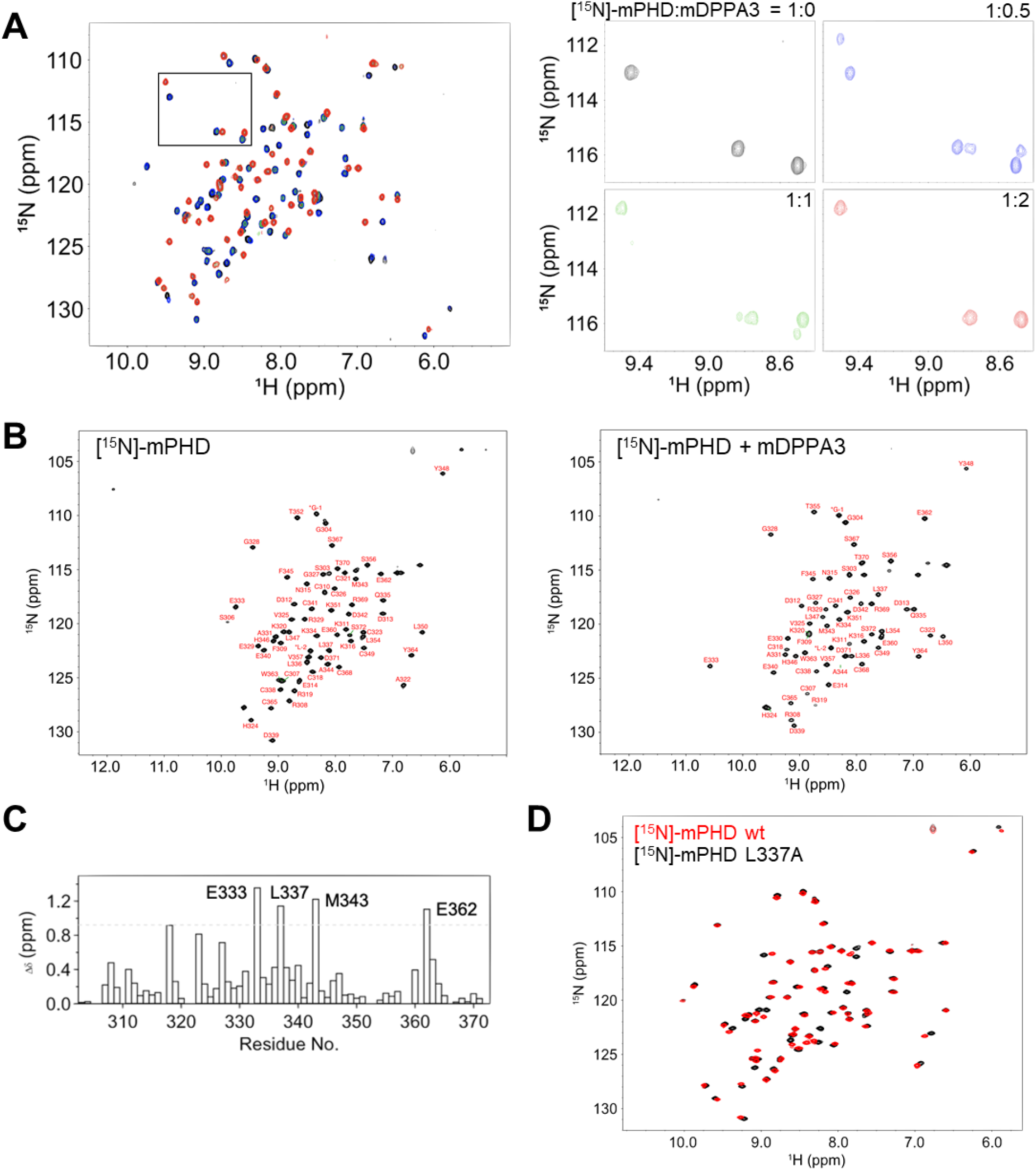
NMR analysis. **(A)** Overlay of ^1^H-^15^N HSQC spectra of 30 μM mPHD showing chemical shift changes upon titration with mDPPA3_76–128W_ of 0 μM (black), 15 μM (blue), 30 μM (green) and 60 μM (red). Square regions inside the HSQC spectra were expanded (right panels). **(B)** Signal assignments in ^1^H-^15^N HSQC spectra of mPHD in the free state (left) and in the complex state with mDPPA3_76–128W_ (right). **(C)** Weighted average chemical shift differences of ^1^H and ^15^N resonances between free mPHD and mPHD in complex withmDPPA3_76–128W_. Dashed line represents mean plus 2 standard deviations. **(D)** Overlay of ^1^H-^15^N HSQC spectra of mPHD wild type (Black) and L337A mutant (red).

**Supplementary Figure S4:**
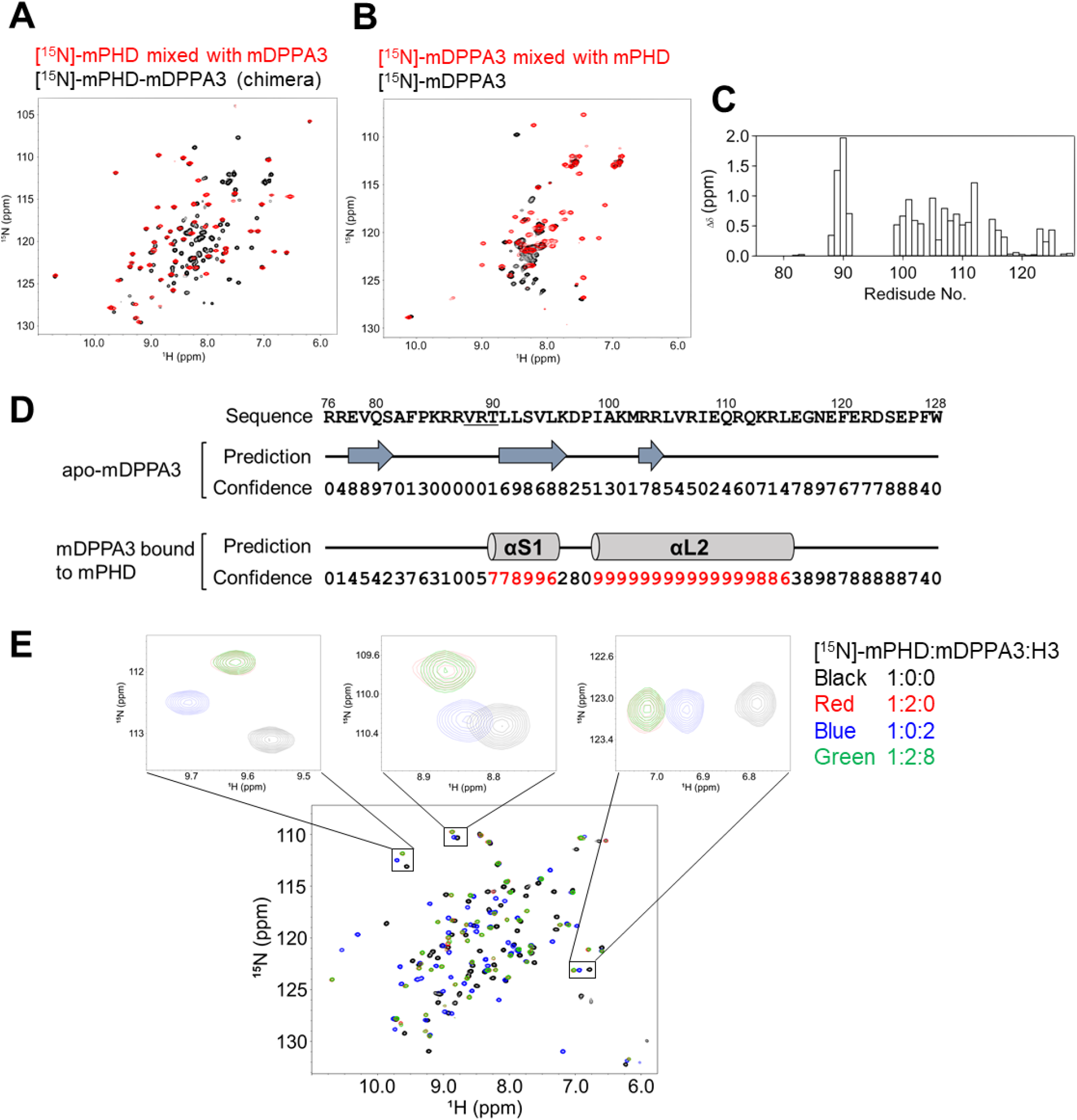
NMR analysis. **(A)** Overlay of ^1^H-^15^N HSQC spectra of [^15^N]-mPHD-mDPPA3 (Black) and [^15^N]-mPHD mixed with mDPPA3_76–128W_ (red). **(B)** Overlay of ^1^H-^15^N HSQC spectra of [^15^N]-mDPPA3_76– 128W_ in the free state (black) and in the complex with mPHD (red). **(C)** Weighted average chemical shift differences of ^1^H and ^15^N resonances between free mDPPA3_76–128W_ and mDPPA3_76–128W_ complex with mPHD. **(D)** Secondary structure prediction using the TALOS-N program for mDPPA3 in the free state and in the complex with mPHD. **(E)** Overlay of ^1^H- ^15^N HSQC spectra of [^15^N]-mPHD in the presence of mDPPA3_76–128W_ and/or the H3 1–37W peptide at molar ratios (mPHD:mDPPA3:H3) of 1:0:0 (black), 1:2:0 (red), 1:0:2 (blue) and 1:2:8 (green).

**Supplementary Figure S5:**
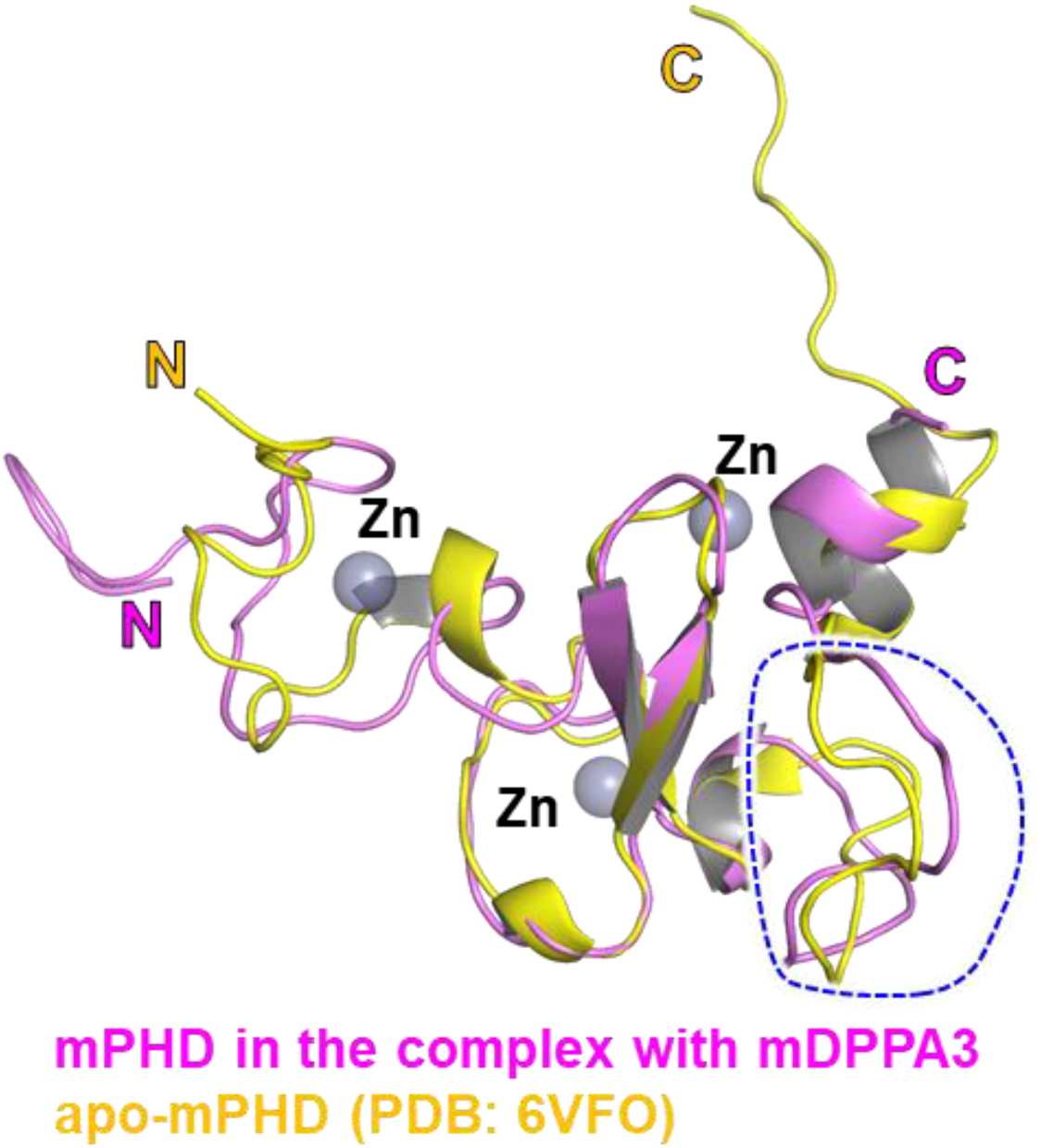
Structural comparison of mPHD and mDPPA3. Overlay of mPHD moiety (colored pink, residues 304–372) in the complex with mDPPA3 on apo-mPHD (PDB: 6VFO, colored yellow, residues 303–380) solved by NMR. Blue dotted line indicates the loop that recognizes N-terminus of H3/PAF15.

**Supplementary Figure S6:**
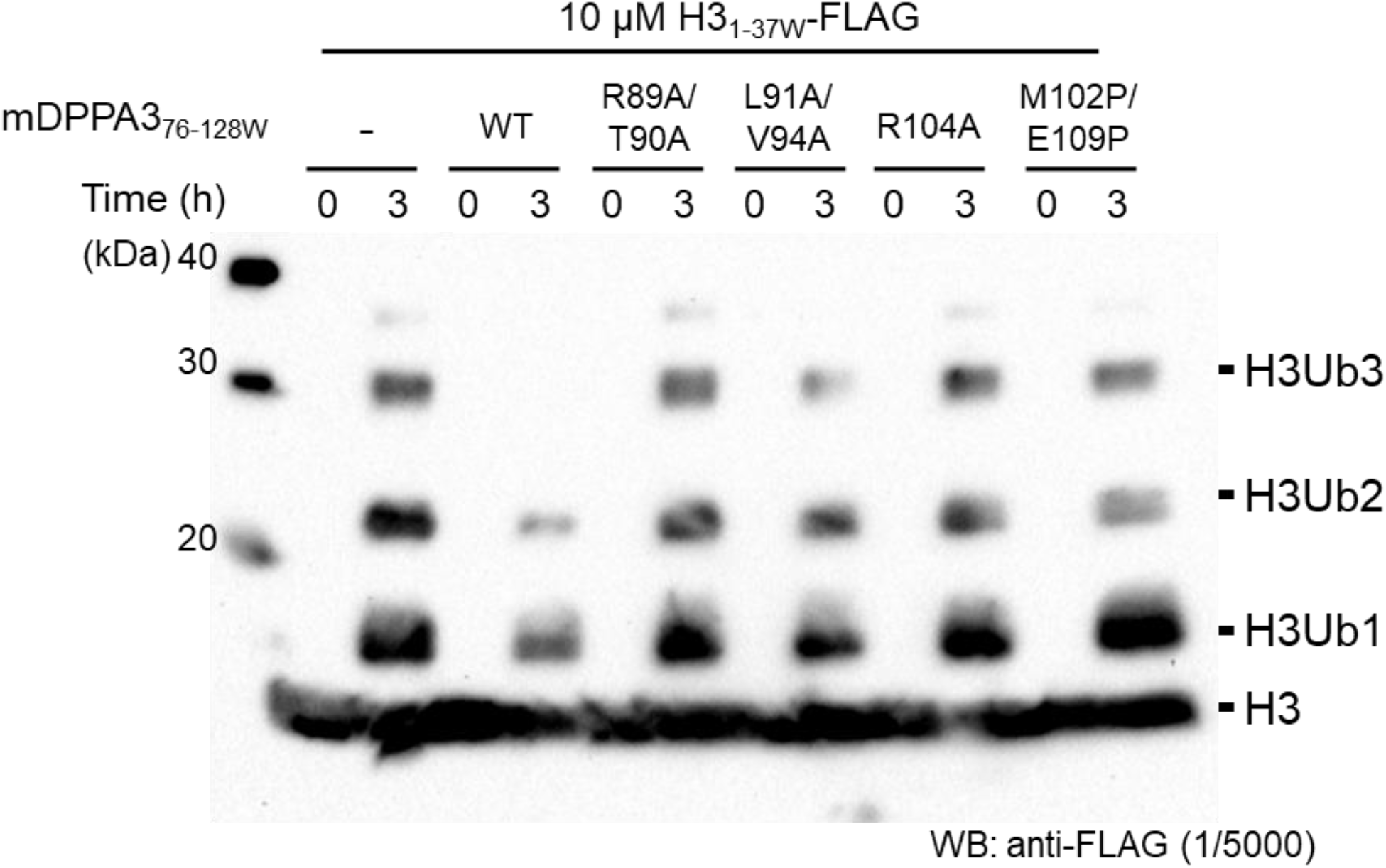
*In vitro* ubiquitination assay of H3 using UHRF1 as a E3-ligase. C-terminal FLAG tagged-H3_1–37W_ was ubiquitinated using in house purified E1, E2, and UHRF1 (E3). The ubiquitinated H3 was detected using anti-FLAG antibody. Equimolar excess of mDPPA3 WT or mutant to H3 was added.

**Supplementary Figure S7.**
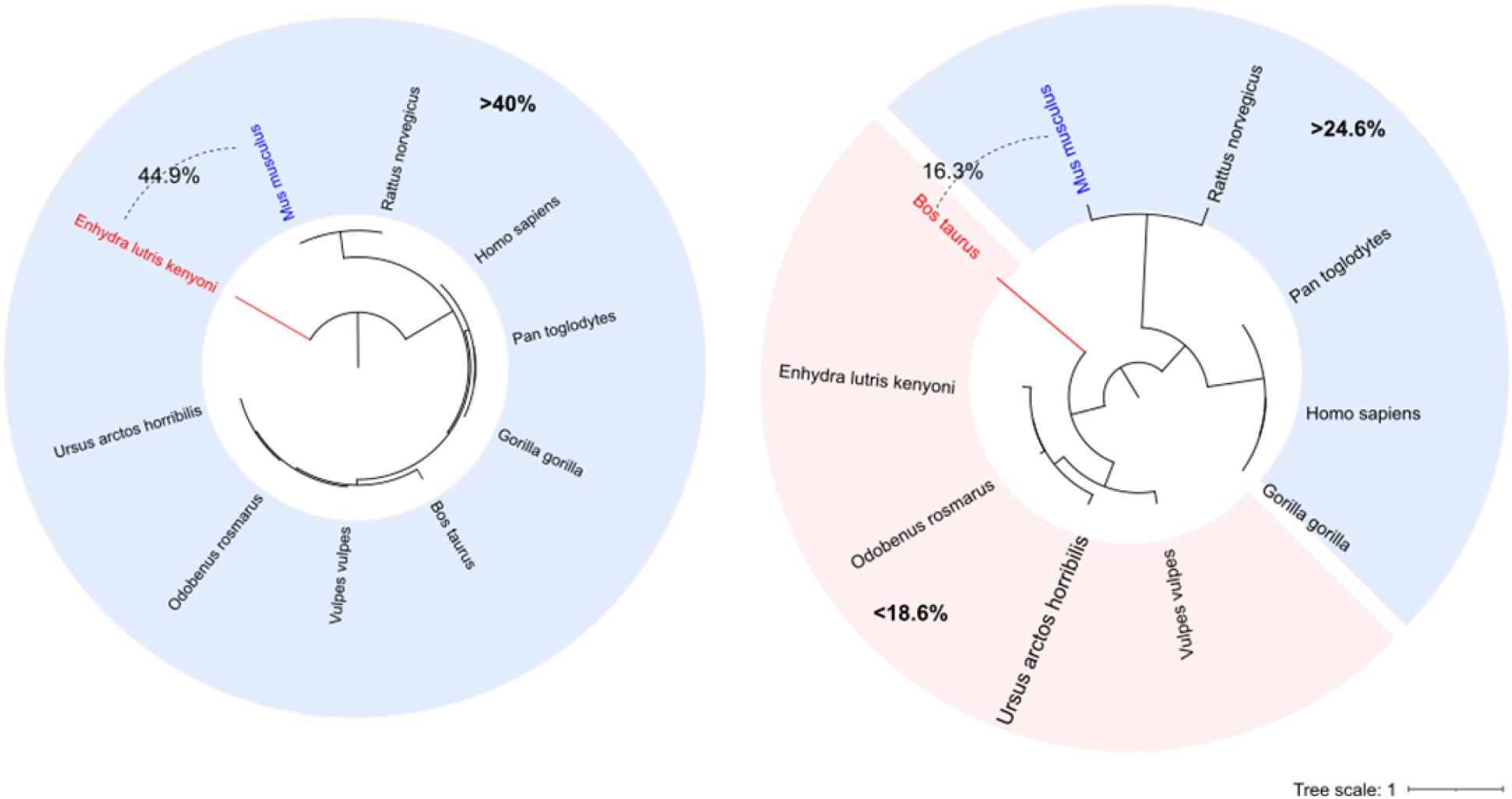
Phylogenetic tree of the UHRF1 and DPPA3. Phylogenetic tree of UHRF1 and DPPA3 in different species rooting by midpoint. The pairwise identities of UHRF1 and DPPA3 between mouse and different species were indicated, light pink (low) and light blue (high). UHRF1 is conserved between species, but DPPA3 is not and forms two clusters, relative conserved (>24.6%) and not conserved (<18.6%).

